# Rapid vaccine induction of macaque HIV-1 V2 Apex broadly neutralizing antibodies with immunogenetic signatures that are potentially translatable to humans

**DOI:** 10.64898/2026.05.27.728024

**Authors:** Severin Coleon, Rasangi Pathirage, Wenge Ding, Elizabeth Van Itallie, Katayoun Mansouri, Lorie Marchitto, Derek W. Cain, Xiaozhi Lu, Varsha Kumari, Laura L. Sutherland, Robert Parks, Andrew Foulger, Thingochuong Le, Lena Smith, Joena Bal, Esther Lee, Alecia Brown, Fangping Cai, Savanna A. Touré, Rebecca Williams, Madison Berry, Ashwin Skelly, Edward F. Kreider, Amie Albertus, Weimin Liu, Houping Ni, Li Zhu, Rumi Habib, Kushal Gandhi, Sampa Santra, Ying Tam, Christopher Barbosa, Drew Weissman, Bette T. Korber, George M. Shaw, Kevin Wiehe, Robert J. Edwards, Kshitij Wagh, Beatrice H. Hahn, Priyamvada Acharya, Barton F. Haynes, Kevin O. Saunders

**Author notes:** Authors contributed equally (Severin Coleon, Rasangi Pathirage).

## Abstract

Due to the complex biology of HIV-1 broadly neutralizing antibodies (bnAbs), HIV-1 vaccines must target unmutated germline precursor antibodies with B cell receptors that have the appropriate specificity and genetic features to develop neutralization breadth. Previous work has shown that immunization or infection in macaques primes unmutated precursor antibodies that target the broadly neutralizing epitope composed of the second variable region and proximal glycans (termed V2-apex site) of HIV-1 envelope (Env). Macaque V2-apex bnAb precursors elicited thus far have homogeneous usage of the second reading frame of a diversity (D) gene found only in macaques, calling into question whether HIV-1 Env immunogens can elicit only this canonical, macaque-specific V2-apex antibody response, limiting human vaccination success. Here, we show that vaccination of rhesus macaques with Env conjugated to a ferritin nanoparticle (OPT4-scNP) rapidly elicited V2-apex bnAbs with distinct D gene segment usage and motifs that are not restricted only to macaques. After two priming immunizations, V2-apex antibodies with non-canonical HCDR3s exhibited greater neutralization potency and breadth than V2-apex antibodies with previously observed HCDR3 motifs. Following sequential boosting immunizations, a V2-apex bnAb lineage with a novel HCDR3-encoded EDGED motif increased its neutralization breadth, including neutralization of viral isolates bearing the N130 glycan, which shields the V2-apex from recognition. High-resolution structures of two members of this V2-apex bnAb lineage, called DH2050, defined a novel binding mode in which it used its EDGED motif to contact the C-strand peptide and the N160 glycan within the V2-apex site. Altogether, we define a novel bnAb binding mode to the V2 Apex and demonstrate that HIV-1 Env sequential vaccination elicits V2-apex bnAbs with immunogenetic signatures that are potentially translatable to the human immune system.

## INTRODUCTION

A protective HIV-1 vaccine could substantially reduce the 1.2 million new HIV-1 infections that happen worldwide each year, including the 32,000 new HIV-1 infections that occur annually in the United States (1). Broadly neutralizing antibodies (bnAbs) are capable of neutralizing genetically diverse HIV-1 isolates and are therefore a major goal of vaccine-induced immunity (2–5). One of the six major sites targeted by HIV-1 bnAbs is the V2-apex(4, 6–8). The V2-apex epitope is located within the part of the HIV-1 envelope glycoprotein (Env) that is most distal from the virus membrane, and is composed of peptide and glycan elements within the first and second variable regions (8–12). V2-apex bnAbs are among the most potent known HIV-1 antibodies, with some antibodies exhibiting IC50 values as low as 0.001 μg/mL across diverse virus panels (4, 13–18). Passive administration of V2-apex bnAbs protect nonhuman primates from mucosal simian-human immunodeficiency virus (SHIV) challenge (3, 19, 20). Notably, approximately 20% of individuals who develop HIV-1 bnAbs generate V2-apex-directed responses, suggesting this class may be reproducibly elicited through vaccination (21, 22).

Despite significant advances in vaccine design, induction of durable, high-titer bnAbs by vaccination in humans has not yet been achieved (6, 23). Successful vaccine induction of bnAbs is thought to require priming with immunogens that engage the unmutated common ancestors (UCAs), or germline precursors, of desired antibody lineages (24, 25). For V2-apex bnAbs, precursor engagement is considered a major challenge because of the rarity of these naïve precursor B cells—estimated at frequencies of approximately 2-200 per ten million naïve human B cells (26–28). Nonetheless, natural HIV-1 Envs such as Q23.17, CM244, WITO, T250-4, and ZM233, and SIV Envs MT145 and CAM13 have demonstrated the capacity to engage some V2-apex bnAb precursors (9, 15, 29–32). Vaccination with engineered Envs based on these natural Envs or these natural Envs themselves have successfully primed V2-apex bnAb precursor antibodies in humanized knock-in mice (30, 31, 33, 34), followed by V2 apex antibody induction in wildtype small animal models (35), and finally in nonhuman primate models (30, 36–40). These previous studies have shown that vaccination primed V2-apex-targeting antibodies that lacked neutralization entirely (38) or lacked neutralization of viruses containing a glycan at position N130 (37). Therefore, overcoming N130 glycan restriction remains a major objective in V2-apex vaccine design (36–39).

Previously, we used neutralization signatures to design a multi-lineage V2 apex bnAb germline targeting Env called CAP256.wk34.c80.OPT4 (OPT4) that when expressed on the surface of a SHIV initiated V2 apex bnAb precursors 400-fold more frequently than wildtype Envs (41). Soluble OPT4 Env trimers were sortase A conjugated to the surface of ferritin nanoparticles (OPT4-scNP) to improve their avidity for V2-apex bnAb precursors to generate an OPT4-based immunogen. OPT4-scNP has been shown to bind to precursors of 14 distinct V2-apex bnAb lineages (41). Two OPT4-scNP primes, followed by an immunofocusing boost with OPT4 containing an additional N-linked glycan at position 187 (termed OPT4-shielded, or OPT4-S) administered as a lipid nanoparticle encapsulated mRNA (mRNA-LNP), elicited serum V2-apex neutralizing antibody responses in all NHPs(41). Whether these V2 apex targeted antibodies could be further affinity-matured to achieve substantial neutralization breadth and potency, and what immunogenetic and structural features such bnAbs might exhibit, remained open questions that we address in the present study.

V2-apex bnAbs share structural and genetic characteristics that facilitate recognition of the trimer apex (9, 29). These antibodies typically possess unusually long (exceeding 20 amino acids) heavy-chain third complementarity-determining regions (HCDR3s) (42, 43), which are frequently anionic and adopt characteristic hammerhead (12, 44, 45), needle (10, 11), or conformations (9). In humans, V2-apex antibodies often utilize YYD motifs (29, 44), whereas structural and immunogenetic analyses of macaque V2-apex antibodies have identified recurrent usage of the IGHD3-15*01 (KIMDB nomenclature) gene segment, which encodes a critical motif in its second reading frame (42, 43, 46). The DDY motif includes sulfated tyrosines that mediate required interactions with Env (43). Because IGHD3-15*01 lacks a direct human ortholog, important translational concerns have been raised about whether vaccine strategies successful in macaques will efficiently elicit comparable V2-apex responses in humans. Accordingly, a major goal of current HIV vaccine development is the design of immunogens capable of inducing V2-apex bnAbs that utilize gene segments and developmental pathways more representative of the human antibody repertoire.

Here, we sought to investigate the developmental pathways of OPT4 Env-primed V2-apex antibodies as they mature into HIV-1 broadly neutralizing antibodies. We primed V2-apex responses with OPT4-scNP immunization and subsequently boosted these responses with a series of HIV-1 envelope immunizations. During this sequential vaccination of macaques, we profiled the immunogenetics, functions, and structures of V2 apex antibodies. Notably, sequential immunization primed a V2-apex bnAb lineage derived from a D gene segment other than IGHD3-15 lacked tyrosine residues, and possessed a new conserved HCDR3 motif—altogether defining a novel binding mode for V2 apex antibodies. Sequential boosting immunizations affinity-matured serum and monoclonal V2-apex antibodies capable of neutralizing viruses with and without the N-linked glycan at position 130. Collectively, the vaccine-induced antibodies identified here provide evidence that V2-apex immunogens can prime subdominant V2-apex bnAb lineages that are not dependent on macaque-specific IGHD3-15*01 gene segments, providing a strong rationale for the clinical evaluation of this sequential vaccine regimen in Phase I trials HVTN 322 and HVTN 326.

## RESULTS

### Sequential immunization elicits serum V2-apex bnAbs in nonhuman primates

We hypothesized that boosting with heterologous envelopes that bind V2-apex antibodies would affinity-mature V2-apex bnAb lineages initiated by OPT4 and OPT4-S immunizations. The OPT4 Env trimer that binds to V2 apex bnAb precursors with low nanomolar binding affinity, respectively (**Figure 1A**), was displayed on the surface of ferritin nanoparticles (OPT4-scNP) (**Figure S1A**). Five rhesus macaques were immunized twice subcutaneously with 100 μg OPT4-scNP adjuvanted with empty lipid nanoparticles (LNPs) ten weeks apart (**Figure 1B**). At study weeks 27 and 45, macaques were immunized with OPT4-S gp150 followed by C.1080 gp150 encoded by mRNA-LNPs, respectively. Lastly, at week 78, macaques received a subcutaneous immunization with BF1266 Env trimer conjugated to mI3 nanoparticles (**Figure S1B**) followed by BF1266 mRNA-LNP six weeks later (**Figure 1B**). Plasma IgG responses after the first OPT4-scNP immunization showed robust binding to autologous OPT4 trimers in all NHPs (**Figure 1C**). Binding specificity was confirmed using a non-specific antigen, BatCoV-NL1404022 RBD, which showed no reactivity. IgG binding was also observed against OPT4-S, although slightly reduced compared to OPT4, suggesting that a subset of the OPT4 primed antibodies targeted its exposed V2 carboxyterminus. We observed that a second OPT4-scNP immunization reproducibly increased IgG binding to both OPT4 and OPT4-S, as well as binding to heterologous HIV-1.C.1080, HIV-1.BF1266, and SIV.CAM13RRK Env trimers in all NHPs. **(Figures 1C and S2A and B**). This result indicated that OPT4-S immunization had immunofocused B cell responses to antigenic surfaces shared by all five widely divergent HIV/SIV Envs.

**Figure 1.**
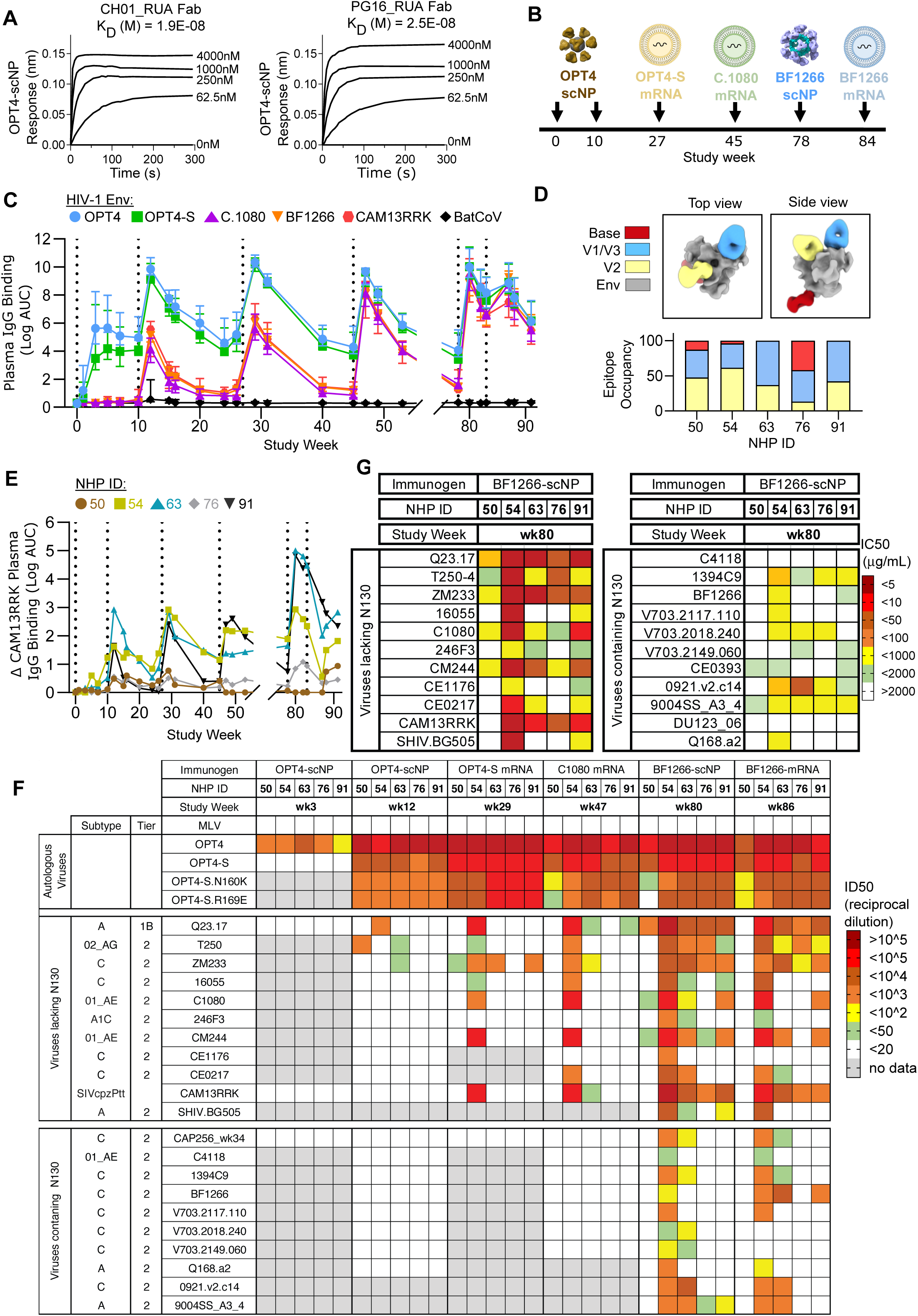
Sequential immunization leads to rapid induction of V2-apex-directed neutralization breadth in NHPs. (A) Biolayer interferometry (BLI) binding kinetics of Fabs derived from human V2-apex bnAb inferred precursors against OPT4 soluble trimers. (B) Immunization regimen of five nonhuman primates. (C) Plasma IgG binding to a panel of soluble envelopes trimers. Binding magnitude is shown as area under the log10 transformed curve (logAUC). (E) EMPEM analysis of Fabs generated from purified plasma IgG isolated at week 15 in complex with OPT4-S soluble trimers. (D) Differential log AUC binding between CAM13RRK and CAM13RRK DKO. (F) Heatmap of serum neutralization titers (ID50 as reciprocal serum dilution) against a panel of virus. (G) Heatmap of neutralization titers (IC50 in μg/mL) against a panel of viruses for IgG purified from week 80 plasma.

To further assess V2-apex specificity, plasma IgG binding was evaluated against OPT4-and OPT4-S with substitutions at positions N160K and K169E (OPT4-DKO and OPT4-S-DKO) that prevent V2-apex bnAb and precursor binding (**Figure S1C and S2A**). No significant differences in plasma IgG binding were observed between OPT4 and OPT4-DKO or between OPT4-S and OPT4-S-DKO in rhesus macaque plasma, likely due to IgG binding to epitopes outside the V2 apex site (**Figure S2**). However, 3D reconstructions from negative stain electron microscopy-based polyclonal epitope mapping (ns-EMPEM) of plasma-derived antibody binding fragments (Fabs) complexed with OPT4-S revealed antibody binding to the V2-apex, in addition to other Env regions, including the base, V1/V3 loops (**Figure 1D**). To better resolve V2-apex–specific responses in plasma, we used CAM13RRK and CAM13RRK DKO (30), which are engineered SIVcpz Env trimers with a V2-apex site similar to OPT4, but the rest of the Env sequence is genetically distant from OPT4. Subtraction of CAM13RRK DKO binding from CAM13RRK binding, revealed the presence of V2-apex–directed antibodies in 3 of 5 NHPs after the second OPT4-scNP immunization (**Figure 1E**). OPT4-S mRNA-LNP immunization increased CAM13RRK binding relative to CAM13RRK DKO at week 29, but only in the three NHPs that already showed differential binding after the second OPT4-scNP immunization. Notably, NHP 54 also showed differential plasma IgG binding between C.1080 and C.1080 DKO (**Figure S2**), indicating the presence of V2-apex antibodies that could recognize the divergent V2 peptide present in C.1080 (**Figures S1D and E**). Because NHP54 showed binding but not serum neutralization against the C.1080 pseudovirus (**Figure 1F, S2C, and S3A**), we hypothesized that C.1080 immunization would affinity mature binding IgG to acquire neutralization activity. A single immunization with C.1080 mRNA-LNP increased plasma IgG binding to C.1080 and BF1266 Env soluble trimers in all animals. NHP54, which had detectable neutralization breadth after OPT4-S mRNA-LNP immunization, exhibited a marked increase in breadth against 11/22 tested viruses (**Figures 1F and S3A**). NHP 91 developed low but detectable heterologous serum neutralization against Q23.17, while NHP63 developed neutralization activity against Q23.17, ZM233, and CAM13.RRK. Thus, boosting with C.1080 improved heterologous neutralization in 3 of 5 macaques (**Figures 1F and S3A**).

To overcome the N130 glycan, which is a significant roadblock for V2-apex bnAb maturation (4, 37), NHPs were immunized sequentially with BF1266 stabilized trimers conjugated to mi3 nanoparticles (BF1266-mi3NP), followed by BF1266 mRNA. BF1266 envelope bears the N130 glycan. A single BF1266-mi3 NP immunization increased serum neutralization breadth in all five NHPs (**Figures 1F and S3A**). NHP 63 and NHP 54 were the strongest responders, achieving neutralization breadth against 18/22 and 22/22 viruses, respectively, including multiple N130-glycan–containing isolates. NHP 76 and NHP 91, which had low serum neutralization breadth, developed responses against 8/22 and 10/22 viruses, respectively. Among the neutralized viruses was the N130-glycan–bearing isolate 9004SS.A3.4. NHP50 was the weakest responder, with detectable serum neutralization against only 5/22 viruses (OPT4, OPT4-S, C.1080, Q23, CM244). To increase the sensitivity for detecting low-titer neutralization, we purified IgG from serum and tested it for neutralization. Purified IgG collected after BF1266-mi3 NP immunization showed broader neutralization and higher potency as compared to the serum of NHPs (**Figures 1F, 1G, and S3**). Purified IgG neutralization of at least 2 viruses with N130 glycans on Env was detected in all NHPs (**Figures 1G and S3B**). After BF1266 mRNA-LNP immunization, heterologous neutralization was still evident, but this immunization predominantly boosted autologous BF1266 neutralization in 4 of 5 NHPs (**Figures 1F and S3**). Together, these results demonstrate that primed V2-apex precursor responses in rhesus macaques following OPT4-scNP immunization can be affinity matured through sequential boosting with immunogens incorporating C-strand diversity and the N130 glycan.

### V2-apex B cell priming and clonal expansion

Env-specific B cells were isolated from inguinal lymph node (iLN) fine needle aspirates (FNA) and peripheral blood mononuclear cells (PBMCs) using fluorescently labeled OPT4, OPT4-S, or BF1266 Env tetramers. FNA sampling of iLN was performed longitudinally at study weeks 3, 15, 24, 31, 49, 82, and 86, and in PBMCs at weeks 12, 53, and 86 (**Figure 2A and S4**). From FNAs, we recovered Env-specific B cells at all weeks except week 49, due to low cell yield. On average, lymph node Env+ B cells were low after a single immunization but were increased relative to week 3 at all subsequent time points (**Figure 2A**). Env-specific B cells were also present in the PBMCs at each time point examined (**Figure 2A**). The phenotype of Env^+^ CD20^+^ B cells was largely class-switched (IgD) regardless of tissue source, whereas total B cells were primarily CD38/IgD naïve-like B cells (**Figures 2B and S5**). iLN-derived Env-specific B cells showed an enrichment of CD38 /IgD cells (**Figure 2B and S5**). On average across the time points sampled, approximately 30% of Env-specific CD20^+^ B cells in iLN were CD38 /IgD (**Figure 2B, Figure S5**). This B cell compartment is important since macaque germinal center B cells are CD20+/CD38 /IgD following germinal center–associated proliferation (47–49).

**Figure 2.**
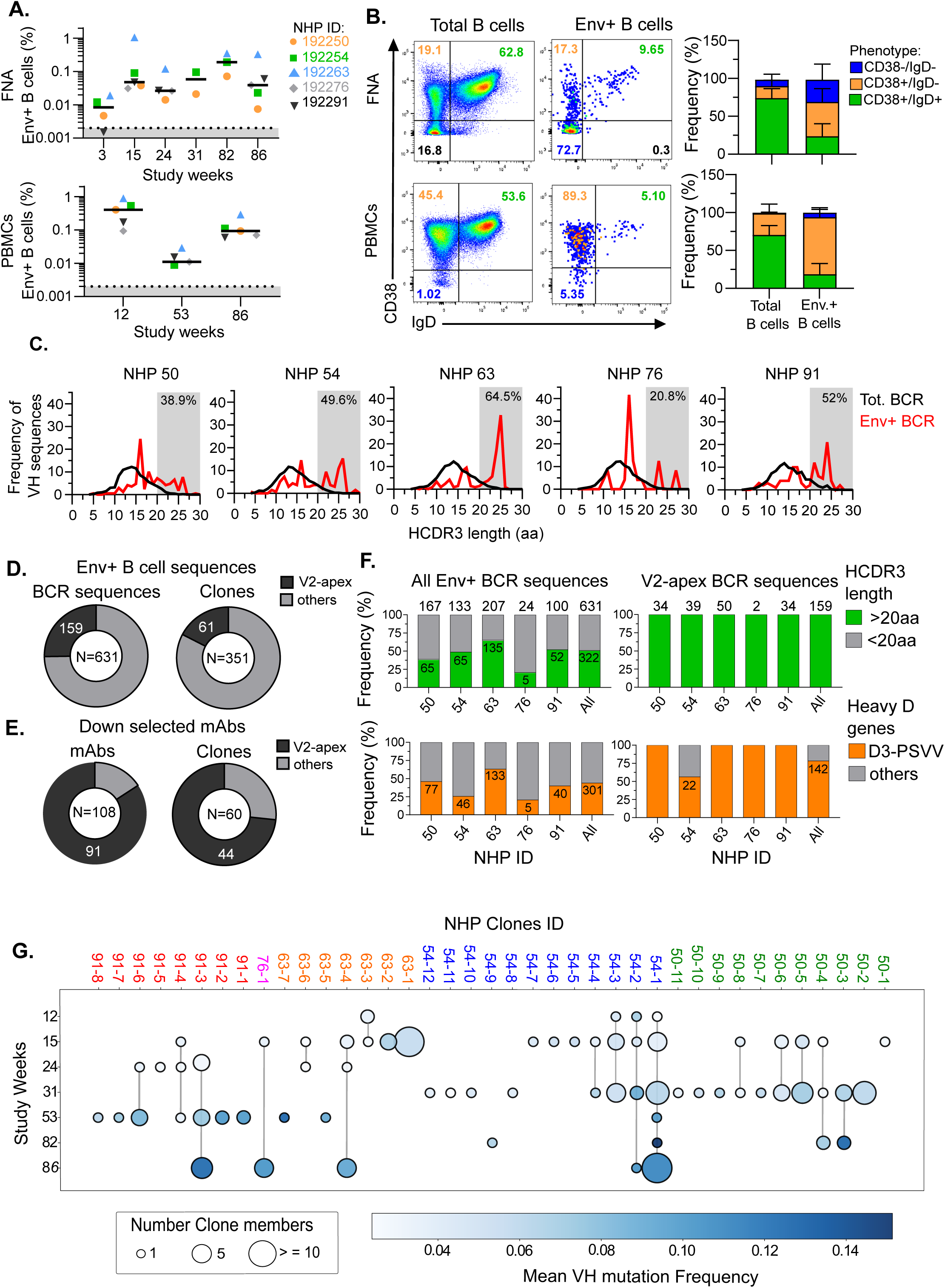
Sequential immunization successfully primed V2-apex B cells precursors in NHPs. (A) Frequency of antigen specific B cells collected in iLN FNA and PBMCs. (B) Representative dot plot and population frequency of antigen specific and total B cells from iLN FNA or PBMCs based on their CD38 and IgD expression. (C) HCDR3 length distribution of sorted antigen specific antibody sequence (red) compared to their B cell immunoglobulin repertoire (black). Gray shaded boxes represent HCDR3 length >20 aa. (D) Proportion of V2-apex directed antibody sequences and their clonal lineages in all Env+ B cells sequences. (E) Proportion of V2-apex directed mAbs and their clonal lineages in down selected mAbs. (F) Frequency of all Env+ BCR sequences and identified V2-apex BCR sequences exhibiting HCDR3 length of at least 20 amino acid and using D3-PSVV gene (Corey Watson Database). (G) V2-apex clonal lineage maturation across the study.

To characterize the Env-specific antibody repertoire primed by OPT4-scNP, we compared the Env-specific antibody sequences isolated from each monkey after two immunizations to their corresponding immunoglobulin repertoire from total B cells. Env-specific antibody sequences were enriched for long (>20 amino acids) heavy chain complementarity-determining region 3 (HCDR3), a hallmark feature of V2-apex bnAb lineages (**Figure 2C**). This enrichment was observed across all animals, but its magnitude varied. NHP 63 exhibited the highest proportion of long HCDR3 Env-specific sequences (64.5%), including an expanded 59-member V2-apex antibody clonal lineage detected in week 15 iLN samples (**Figure S6**). In contrast, NHP 50 and NHP 76 had an enrichment of shorter (∼16 amino acid) HCDR3 sequences with relatively fewer long HCDR3 Env-specific antibodies (38.9% and 20.8%, respectively) compared to other NHPs (**Figure 2C**).

To determine the specificity and function of the Env-specific B cells isolated from all six FNAs and three PBMC samples, we amplified heavy and light chain sequences by RT-PCR and cloned them into expression vectors. Recombinant antibodies were expressed, and three days post transfection cell-free supernatants containing the antibodies were screened by ELISA against a panel of HIV-1 Env trimers (**Figure S4A**). Antibodies exhibiting strong binding to OPT4 and OPT4-S, reduced binding to OPT4-S DKO, and or cross-reactivity to heterologous Env proteins were down-selected for monoclonal antibody (mAbs) production and further characterization (**Figure S7**). One hundred eight out of 631 isolated BCR sequences were down-selected for mAb production across all NHPs (**Figure 2D and E**). Down-selected mAbs were characterized by virus neutralization assays (**Figures S8 and S9**), negative-stain electron microscopy (NSEM) (**Figure S10**), and ELISA binding assays (**Figure S10**), with 91 out of 108 down-selected mAbs confirmed to be V2-apex specific (**Figure 2E**). Eight-six percent of the 91 antibodies exhibited OPT4-S neutralization activity (**Figure S9**). Overall, 159 BCR sequences, representing 25% of the total Env-specific sequences, were determined to be V2 apex specific (**Figure 2D**). Across all animals, 51% (322/631) of Env-specific antibody sequences exhibited long HCDR3 (≥20 amino acids), and 47.7% (301/631) were derived from IGHD3-PSVV*01 (equivalent to IGHD3-15 in the KIMDB nomenclature) (**Figure 2F**). As expected, the confirmed V2-apex BCR sequences exhibited long HCDR3. They also showed the exclusive usage of the IGD3-PSVV gene in all macaques except NHP 54, where only 57% of its confirmed V2-apex BCR sequences used IGD3-PSVV (**Figure 2F**). Longitudinal analysis of 39 V2-apex clonal lineages across all NHPs revealed the isolation of 20 expanded lineages, with 17 lineages isolated at two timepoints or more. Interestingly, all macaques except NHP 76 showed the engagement of at least six distinct V2-apex clonal lineages, with somatic hypermutation increasing over time (**Figure 2G**). Consistent with increased somatic mutation over time, the lineages showed increased neutralization potency against OPT4 and OPT4-S, and the development of neutralization against heterologous viruses bearing the N130 glycan (**Figures S8 and S9**).

### OPT4-scNP immunization primed stereotypical and atypical V2-apex B cells

The binding angle of approach for 31 mAbs isolated from weeks 15 and 24 iLN samples was characterized by NSEM analysis. Three distinct binding approach angles to the V2-apex region were observed for the antigen-binding fragment (Fab) of the 31 mAbs in complex with OPT4-S (**Figure 3B and S10**). These three binding approach angles were distinct from human V2-apex bnAb precursors (PG9_RUA and CH01_RUA) and rhesus macaque V2-apex precursor 41238 UCA (43) (**Figure 3B**). One group of mAbs, called the DDY mAbs, exhibited similar binding approach angles and had immunogenetic features stereotypical of rhesus macaque V2-apex antibodies, including long HCDR3 loops (>20 amino acids), an overabundance of HCDR3 tyrosines, IGHD3-PSVV*01 gene usage, and a D gene-encoded aspartate-aspartate-tyrosine (DDY) HCDR3 motif (36, 37, 42, 43, 46) (**Figure 3C and S10**). OPT4-scNP consistently primed stereotypical ‘DDY’ V2-apex precursors across all immunized animals (**Figure 3C**).

**Figure 3.**
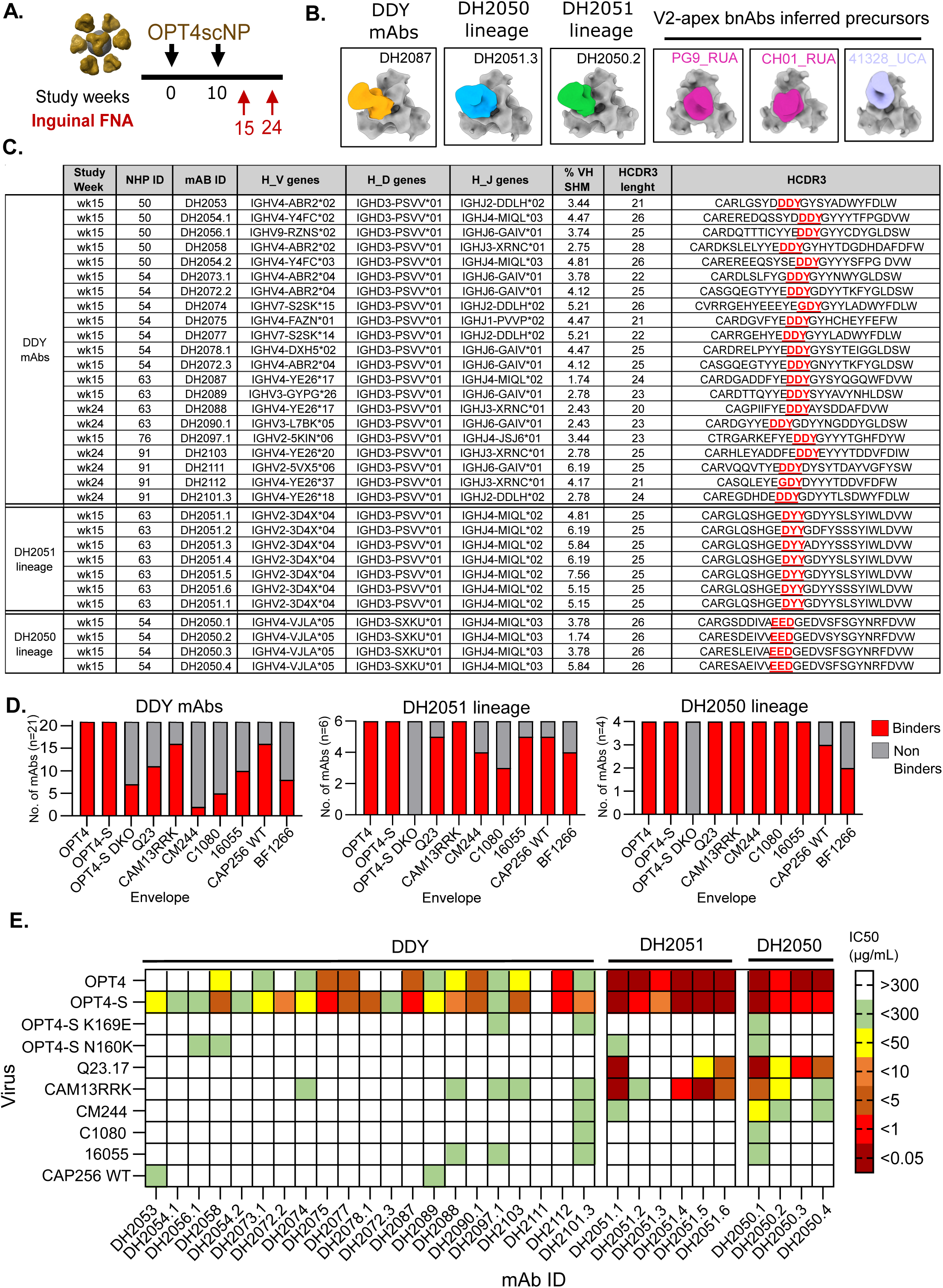
Immunization with OPT4-scNP primed stereotypical and atypical V2-apex B cells precursors. (A) Immunization regimen with OPT4-scNP and iLN FNA collection time point. (B) NSEM representative of isolated V2-apex mAbs complexed with OPT4-S soluble trimers and V2-apex bnAb inferred precursors complexed with OPT4. (C) Immunogenic table of V2-apex antibodies isolated following 2 immunization with OPT4-scNP. (D) Bar graph representation of V2-apex antibodies isolated following 2 immunization with OPT4-scNP, that bind or not to a panel of soluble envelopes trimers. Antibodies with binding log AUC values higher than the negative control antibody log AUC + 3 standard deviations were considered positive. (E) Neutralization IC50 of V2-apex antibodies isolated after 2 immunizations with OPT4-scNP against a panel of virus.

The remaining 10 antibodies comprised two expanded clonal lineages in NHP 54 (DH2050) and NHP63 (DH2051) exhibited alternative structural and genetic features compared to the DDY mAbs. Antibody clone DH2051 was derived from IGHD3-PSVV*01, but lacked the DDY motif and instead had an aspartate-tyrosine-tyrosine (DYY) motif (**Figure 3C and S11**). Antibody clone DH2050 was derived from non-canonical D gene IGHD3-SXKU*01, had a glutamate-aspartate-glycine-glutamate-aspartate (EDGED) HCDR3 motif spanning positions 100c to 100g and only one or two tyrosines in its HCDR3 (**Figure 3C and S11**). When tested for binding by ELISA against a panel of soluble Env trimers, stereotypical ‘DDY’ and atypical mAbs showed strong binding to OPT4 and OPT4-S, with reduced binding to OPT4-DKO (**Figure 3D and S10C**). Stereotypical ‘DDY’ V2-apex mAbs and atypical V2-apex mAbs showed binding breadth, which included binding to two N130-glycan–bearing envelopes CAP256.wk34 and BF1266 (**Figures 3D and S10C**). The three antibody types differed primarily in the proportion of antibodies within each type or lineage that was capable of binding CM244 and C.1080 Envs. Most atypical V2-apex lineages reacted with CM244 (4/6 DH2051; 4/4 DH2050) and C.1080 (3/6 DH2051; 6/6 DH250), whereas only a small fraction of the DDY mAbs bound these two Envs. Similarly, neutralization activity of the stereotypical V2 apex antibodies was less potent and less broad than that of the atypical antibodies (**Figure 3E**). Seven of ten atypical V2 apex antibodies neutralized at least one heterologous virus with an IC50 below 10μg/mL (**Figure 3E**). Altogether, OPT4-scNP engaged stereotypical V2-apex precursors in all macaques, but also engaged more rare, atypical V2-apex bnAb precursors that showed heterologous neutralization after only two immunizations.

### Maturation of atypical V2-apex antibodies to acquire neutralization breadth

The vaccine-induced atypical antibody clone DH2050 persisted throughout the vaccine regimen (**Clone 54-1 in Figure 2G**). We successfully recovered 26 BCR sequences from the DH2050 clonal lineage across five different time points. Phylogenetic reconstruction from these 26 BCR sequences showed that the antibody sequences identified after the BF1266 boost had diverged the most from the unmutated common ancestor (**Figure 4A**). Further analysis of the antibody sequences revealed that the EDGED motif was unchanged in 22 of 26 clonal members and was part of a long N2-region rather than the germline D gene segment (**Figure 4B and S11**). Additionally, a 4-amino-acid insertion in the HCDR2 was observed in sequences isolated after the OPT4-S mRNA-LNP immunization. Fab binding affinities were measured throughout maturation of the DH2050 clone. Fabs selected from weeks 15, 31, 53, and 86 showed a staged progression for binding to vaccine immunogens (**Figure 4C**). All tested Fabs had nanomolar affinity for OPT4-S regardless of when they were isolated (**Figure 4C**). In contrast, Fabs with detectable affinity for heterologous boost Envs, C.1080 and BF1266, were first detected after the OPT4-S mRNA-LNP boost at week 27 (**Figure 4C**). Fab affinity to either heterologous Env peaked after the Env was given as an immunogen (**Figure 4C**). DH2050.11 and DH2050.15 Fabs had the highest affinities for C.1080 and BF1266 Envs respectively. The variable regions of DH2050.11 and DH2050.15 included the HCDR2 insertion. Similarly, there was enhanced neutralization potency from lineage members carrying the HCDR2 insertion (**Figure 4B)**, suggesting that both HCDR3 and HCDR2 contributed to the functional breadth of this clonal lineage (**Figures 4D and 4E**). Interestingly, a clear relationship between heavy chain somatic hypermutation and neutralization breadth was observed, with early lineage members such as DH2050.2 showing low mutation percentages (∼1.7% VH mutation) and neutralization against autologous viruses only (OPT4 and OPT4-S) (**Figure 4D**). In contrast, clonal members isolated at later time points exhibited somatic VH mutation percentages up to 15.1.% and neutralized all viruses tested (9/9) with IC50 values below 1 µg/mL. We assessed neutralization breadth of the more mutated clonal members against viral isolates bearing the N130 glycan and observed 5 out of 7 clonal members tested showed potent neutralization against N130 viruses (**Figures 4E, S8 and S9D**). We isolated mAbs from weeks 15 to 86 from all NHPs. While clones from NHP 63 such as DH2095 exhibited broad neutralization, antibodies in the DH2050 clone exhibited the broadest neutralization (**Figure S9**).

**Figure 4.**
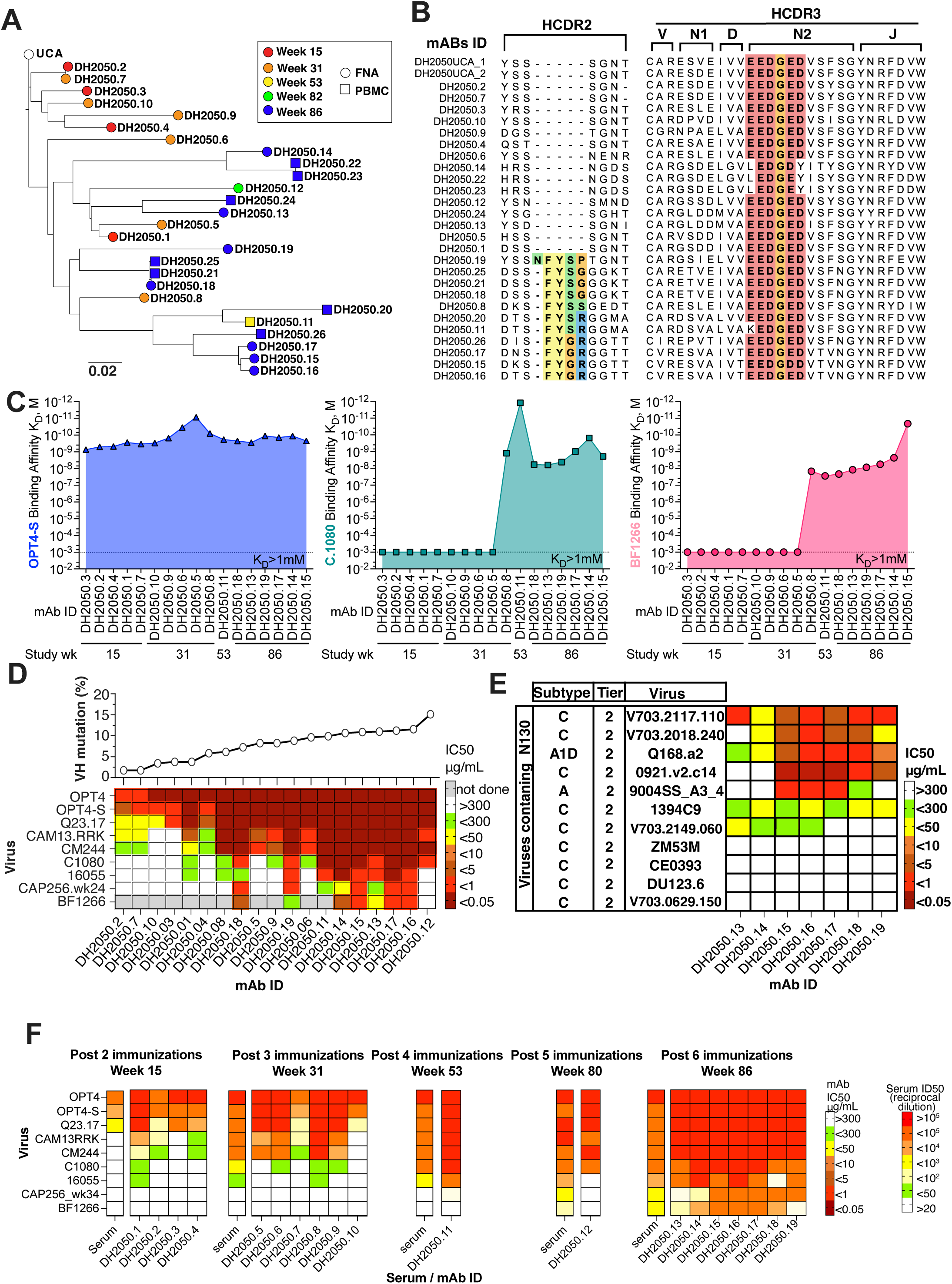
Vaccine-induced atypical V2-apex bnAbs precursors mature and acquire neutralization breadth upon sequential vaccination. (A) Phylogenic tree of DH2050 clonal lineage members isolated during the course of the study. (B) Amino Acid sequence alignment and VDJ mapping of DH2050 clonal members. (C) Binding affinity shown as equilibrium dissociation constant (K_D_) for 17 DH2050 clone antibodies isolated at various time points as indicated. Binding affinity was measured for OPT4-S (left), C1080 (middle), and BF1266 (right) Envs. (D) DH2050 lineage maturation. Top: Mutation frequency of DH2050 lineage members heavy chain variable region (VH). Bottom: Neutralization IC50 of DH250 lineage members against a panel of viruses. (E) DH2050 clonal members neutralization IC50 titers against an extended panel of heterologous envelopes exhibiting N130 glycan. (F) Heatmaps of neutralization IC50 titers for DH2050 clonal members compared to NHP54 serum neutralization ID50 titers following sequential immunization.

Lastly, we compared the neutralization breadth between DH2050 antibodies and NHP 54 serum to explore if this clonal lineage conferred the exceptional serum neutralization breadth observed in NHP 54. Sequential comparison of the neutralization breadth of DH2050 clonal members isolated after each immunization with the breadth detected in NHP 54 serum, revealed stepwise increases in DH2050 lineage and serum neutralization breadth following each immunization (**Figure 4F**). These results demonstrate the engagement and maturation of a V2-apex bnAb lineage through sequential vaccination and its temporal association with the acquisition of serum neutralization breadth.

### DH2050 lineage HCDR3 interactions with the V2 apex

We next determined structures using cryo-electron microscopy (cryo-EM) to visualize details of how the DH2050 lineage antibodies bound to the Env V2 apex. We solved the structures of CM244 Env trimers bound to either the DH2050.1 Fab or the DH2050.8 Fab, with global map resolutions of 3.95 Å and 4.23 Å, respectively (**Figure 5A, 5B, and S12-S15**). The DH2050 lineage antibodies engage the Env apex with an orientation and angle of approach that is reminiscent of the V2-apex bnAb PG9 (**Figure 5A**). Examining the details of the Env/antibody interfaces reveals anionic HCDR3-dominated interactions mediated by DH2050.1 and DH2050.8 with the Env V2 apex (**Figure S16)**. Although the vaccine-elicited antibodies are similar to the V2-apex bnAbs elicited by natural infection in this regard, several key differences were noted. While the bnAbs elicited via natural infection penetrated the Env glycan shield to contact the V2-apex protein surface with their CDRH3 loops (except CH01, for which CDRH1 contacts with the Env protein surface were noted in addition to the HCDR3 contacts), DH2050.1 and DH2050.8 utilized CDRs from both heavy and light chains to contact protein residues at the V2-apex to interact with a larger Env protein surface (**Figure 5C and S16**). The shapes of the bound DH2050.1 and DH2050.8 HCDR3 loops are reminiscent of that of the CH01 antibody, but while the CH01 predominantly contacts the protein surface of one Env protomer (9, 50), the vaccine-elicited DH2050.1 and DH2050.8 antibodies engage the protein surfaces of two protomers.

**Figure 5.**
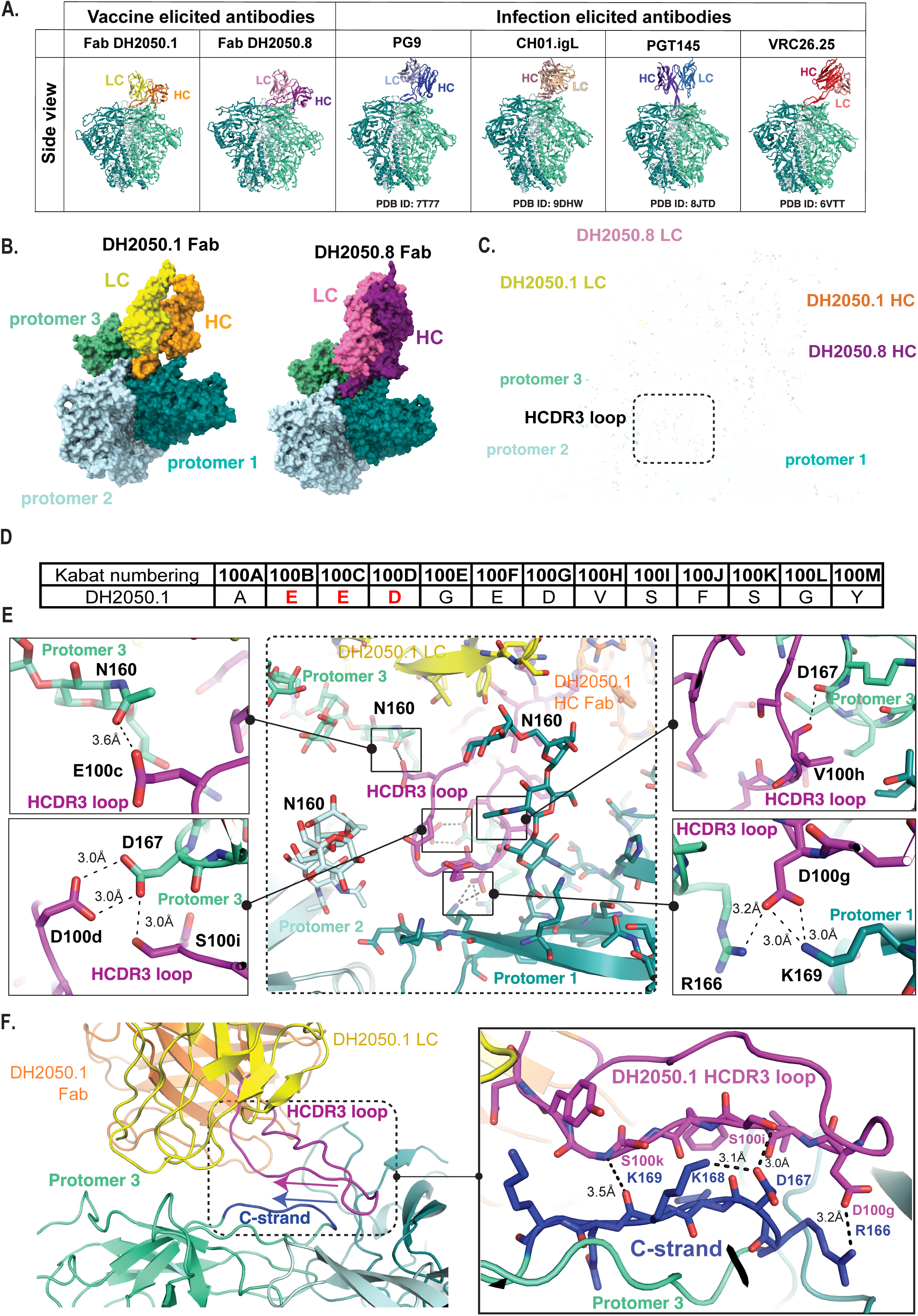
Cryo-EM structures of two DH2050 mAbs reveal a novel V2 apex recognition mode. (A) Cryo-EM maps of DH2050 lineage members DH2050.1 and DH2050.8 Fabs in complex with CM244 compared with cryo-EM model of bnAbs PG9 (PBD:7T77); CH01 (PDB:9DHW); PGT145 (PDB:8JTD) and VRC26.25 (PDB:6VTT). (B) Cryo-EM models of DH2050.1 and DH2050.8 Fabs in complex with CM244. (C). Superposition of DH2050.1 and DH2050.8 Fabs in complex with CM244. (D) Kabat numbering of DH2050.1 HCDR3 amino acid sequence. (E) Hydrogen bonds Interactions between DH2050.1 HCDR3 and CM244 V2-apex C-strand. (F) DH2050.1 Fab interaction with CM244 V2-apex protomers.

Both mAbs exhibited protruding HCDR3s contacting a quaternary epitope at the V2-apex of the envelope (**Figures 5E and S18**). The DH2050.1 Fab HCDR3 engaged the cationic, lysine rich C-strand of one Env V2 protomer in an antiparallel strand-strand interaction. Notably, the serine (S) residues at position 100k and 100i of the HCDR3 make respective interactions with the lysine (K) at position 169 and the highly conserved aspartic acid (D) at position 167 of the C-strand (**Figure 5E and 5F**). This interaction was reminiscent of the VRC26.25 Ser100f interaction with Env D167 (51). Residue E100c within the HCDR3 _100c_EDGED_100g_ motif, interacted with the N160 glycan, while D100d interacted with Env D167, and D100g contacted Env R166, providing a structural explanation for the conservation of the _100c_EDGED_100g_ motif in the DH2050 lineage (**Figure 5E**). The HCDR2 insertion also added contacts with the Env indicating its key role in improving binding and neutralization activity during lineage maturation (**Figure S16**). Thus, this novel binding mode was characterized by key contacts by the EDGED motif and HCDR2 insertion in the DH2050 heavy chain.

## DISCUSSION

In this study, vaccination with OPT4 sc-NPs rapidly engaged V2-apex bnAb precursors that persisted and matured over time, resulting in increased serum neutralization breadth in NHPs. OPT4-scNP efficiently primed a polyclonal V2-apex antibody response consisting of V2-apex bnAb precursors with canonical DDY motifs and distinct EDGED or DYY motifs. The DYY motif was notable since human V2 apex bnAbs typically possess di-tyrosine motifs(27). Notably, elicitation of V2-apex antibody lineages with one particular HCDR3 motif did not interfere with induction of a second type of V2 Apex bnAb lineage since both types of bnAb lineages occurred in the same NHP. Moreover, the elicitation of both types of V2-apex antibody lineages in the same NHP was associated with the greatest serum neutralization breadth.

Here we identified DH2050, a vaccine-elicited V2-apex bnAb lineage characterized by a highly conserved EDGED motif that lacks tyrosine residues. Notably, this HCDR3 motif was not germline encoded, but instead arose through nucleotide insertions during V(D)J recombination. These findings demonstrate that germline gene segment-encoded HCDR3 motifs are not an absolute requirement for V2-apex bnAb development, and that n-addition adds to the pool of V2 apex precursors that can be engaged during vaccination. These findings are consistent with observations of Habib and colleagues who found EDDY motifs encoded by non-templated sequences in naïve human B cells (50). The diverse array of V2-apex precursors engaged by OPT4 irrespective of D-gene usage supports its translational potential for humans.

The binding mode of DH2050 identifies a novel V2 apex binding mechanism. Typically, V2 apex bnAbs like PGT145 interact with the apex hole formed by the three protomers of the trimer (10). This type of interaction usually comes from a needle-like, beta-hairpin HCDR3 conformation, which DH2050 does not have. Additionally, V2 apex bnAbs like PG9 and CH03 make extensive contact with the C-strand and have limited HCDR3 insertion into the apex hole(9, 12, 51). The limited apex hole insertion and HCDR3 juxtaposition with the C-strand makes the binding mode of DH2050 antibodies most similar to PG9/CH03. However, the lack of tyrosine sulfation in the HCDR3 of DH2050.1 or DH2050.8 makes these antibodies distinct from PG9(12, 51), VRC26.25 and PGT145, which use sulfated tyrosines to contact C-strand amino acid side chains(10, 12, 51). CH03 and VRC38.01 lack sulfation of their HCDR3 tyrosines but still utilize tyrosines for C-strand contacts(9, 18). In contrast, DH2050.1 and DH2050.8 antibodies utilize Glu, Asp, and Ser amino acids to contact C-strand amino acids that are usually contacted by V2 apex bnAb HCDR3 tyrosines. While we have no structural evidence for tyrosine involvement in binding, it should be noted that six DH2050 lineage members somatically mutated to add a tyrosine in their HCDR3. Thus, the affinity maturation process may eventually select for amino acids typically found in V2 apex bnAb HCRD3s. Nonetheless, the currently observed DH2050 antibodies contact typical sites within the V2 apex, but do so using a unique set of paratope interactions.

Our results demonstrate sustained engagement and continued affinity maturation of 17 vaccine-induced V2-apex lineages for up to 71 weeks. Thus, B cells either reentered or persisted in germinal centers where somatic mutations were introduced and higher affinity B cell receptors were selected. For B cells to remain in the germinal center, the affinity-dependent model of germinal centers posits B cells with intermediate affinity stay longer in the GCs undergoing additional rounds of maturation (52), whereas high affinity B cells migrate out of germinal centers as antibody secreting cells (ASC)(53). We chose envelope immunogens with V2-apex specific binding, but the antibodies lacked neutralization activity against the corresponding virus. In particular, members of the DH2050 lineage were continuously isolated from inguinal lymph nodes throughout the 86 weeks of the study. The sequential vaccine concept relies on the continued somatic mutation of B cell lineages with each boost.

It is debated whether memory B cells can be recruited back into secondary germinal centers (54), with immunological phenomena such as antibody-mediated feedback being proposed to suppress re-entry of antigen binding B cells into secondary germinal centers (55, 56). We observed antibody clones that were present over time, continued to mutate, and improved in affinity. Since previous studies have shown that HIV-specific germinal center B cells can persist for more than 30 weeks post vaccination (57), we cannot distinguish whether B cell lineages including DH2050 persisted in germinal centers or reentered germinal centers upon vaccination. However, we show the timing between immunizations and the immunization site used here are conducive to continued selection of bnAb lineage members.

Finally, the N130 glycan on Env has limited V2 apex antibody neutralization breadth in previous studies, and it has been suggested that the inclusion of immunogens with the N130 glycan will be needed to overcome this roadblock (37). We demonstrate here the capacity of BF1266, an Env with an N130 glycan and diverse C-strand sequence that we chose as a late-stage boost based on Env neutralization signatures of mature V2 apex bnAbs (58), to expand neutralization breadth to overcome the hindrance of the Env N130 glycan. While protein BF1266 Env nanoparticle immunization expanded serum neutralization breadth against multiple N130-bearing viruses, BF1266 mRNA-LNP immunization primarily recalled responses restricted to the BF1266 isolate. These two immunogens differ in that BF1266 mi3-NP presents 20 stabilized Env envelope trimer proteins clustered on the surface of a nanoparticle, suggesting a potential role of antigen avidity in driving humoral responses against N130-bearing viruses. Thus, acquisition of neutralization breadth of viruses with Envs glycosylated at N130 may require immunization with N130-bearing Envs and their presentation in high-avidity multimeric format (59). The applicability of this strategy for eliciting broad and durable V2-apex bnAb responses that overcome the N130 glycan in humans will be determined in phase I trials HVTN322, HVTN 326, and HVTN 809.

## MATERIALS AND METHODS

### Study Design

The primary objective of this study was to determine the immunogenicity of a sequential vaccine aiming to affinity-mature V2-apex bnAbs in macaques. The immunologic endpoints associated with this objective were serum neutralization breadth and potency and monoclonal antibody neutralization breadth and potency. Our secondary objective was to classify V2-apex antibodies elicited by the sequential vaccine. The secondary immunologic endpoints for this objective were antibody V_H_ and V_L_ sequencing of V2-specific B cells and antibody angle of approach to Env. This study included five female Indian-origin rhesus macaques (ID number 192250, 192254, 192263, 192276, 192291) ages 9-11 years old. The rhesus macaques weighed between 6.5 and 9.1 kg. Macaques were housed at Bioqual Inc., Rockville, MD, in accordance with AAALAC guidelines. The study protocol and all veterinarian procedures were approved by the Bioqual IACUC as stated in a memorandum of understanding with the Duke IACUC. All procedures were conducted based on standard operating procedures. Macaques were repurposed from a HIV MPER peptide liposome immunization trial. As part of the MPER liposome peptide immunization study, macaques were administered 450 mcg of MPER DH511 peptide liposome plus Alum at week 24. The total dose of the vaccine was divided in half and administered intramuscularly to each mid-thigh (225 mcg per leg). Prior to and during this present study serum was tested for MPER-specific neutralization and was negative at all timepoints tested. Macaques were administered 100 mcg of OPT4 sortase A conjugated nanoparticle (OPT4-scNP) mixed with 1.9 mg of empty lipid nanoparticle (Acuitas Therapeutics, LLC) at weeks 0 and 10. OPT4-scNP protein and LNP were diluted in sterile saline up to 1 mL and the total dose of the vaccine was divided in half and administered subcutaneously to each mid-thigh (0.5 mL per leg). At week 27, Macaques were administered 200 mcg of CAP256.wk34.V2OPT.R189TmRNA5-TKR-310 FM-3343B (OPT4-S mRNA). The total dose of OPT4-S mRNA vaccine was divided into four injections with 2 doses administered intramuscularly in each mid-thigh and 2 doses administered subcutaneously in each deltoid (50 mcg per site). At week 45, macaques were administered 200 mcg of lipid nanoparticle (Acuitas Therapeutics, LLC) encapsulated C1080.mRNA5_KW-TKR476 FM 3643B (C.1080 mRNA), with the total dose of C.1080 mRNA vaccine divided in 4 with 2 doses administered subcutaneously in each mid-thigh and 2 doses administered intramuscularly in each deltoid (50 mcg per site). At week 78, macaques were immunized with 100 mcg of BF1266.6R.DS.SOSIPv8.4-mi3 nanoparticle (BF1266-mi3NP) mixed with 1.9 mg of empty lipid nanoparticle (Acuitas Therapeutics, LLC). BF1266-mi3NP and LNP were diluted in sterile saline up to 1 mL and the total dose of the vaccine was divided in half and administered subcutaneously to each mid-thigh (0.5 mL per leg). Finally, at week 84, rhesus macaques were administered 200 mcg of lipid nanoparticle encapsulated pUC-ccTEV-BF1266.mRNA5-A101-TKR522 (BF1266 mRNA). The total dose of BF1266 mRNA vaccine was divided in 4 with 2 doses administered subcutaneously in each mid-thigh and 2 doses administered intramuscularly in each deltoid (50 mcg per site). Peripheral blood was collected by venipuncture in EDTA tubes while macaques were sedated per standard operating procedures and shipped from Bioqual to Beth Isreal Deaconess Medical Center by overnight delivery. Fine needle aspirates (FNA) were collected from macaques by palpating the draining inguinal lymph node and sampling cells with a needle. Collected cells were stored in Hanks balanced salt solution (HBSS) media with EDTA in a conical tube and shipped overnight from Bioqual to Duke University, where they were stained for fluorescence activated cell sorting. If FNA cell pellets were red in color, then the cells were washed up to two times with ACK lysis buffer to eliminate red blood cells. PBMCs and inguinal lymph node FNA cells typically showed >80% viability. All macaques were monitored visually for adverse events and blood chemistry and cell counts were assessed throughout the study.

### Design of CAP256.OPT4/OPT4-S Env and SHIV.CAP256.OPT4/OPT4-S

CAP256.OPT4/OPT4-S envelopes were design as previously described (41). Briefly, we designed an HIV-1 Env to include features enabling binding to multiple rhesus and human V2 apex bnAb germline precursors. We replaced V1 and V2 loops from CAP256.wk34.c80 Env sequence with loops from other naturally occurring Envs that exhibited optimal length and charge for sensitivity, in this case ZM233 for V1 and T250 for V2 (60). Next, we made two substitutions in the C-strand K169R and K170R and another one N130H to remove the N-linked glycan at position 130. Finally, additional glycan at position 339, 397 and 413 were implemented. Recombinant soluble protein was made with the RnS2 stabilization approach as described previously (51). A variant of this Env (CAP256.OPT4-S) was also created to introduce a potential N-linked glycan at residue N187 to shield an immunodominant strain-specific epitope in the V2 carboxyterminus. The nucleotide sequences of CAP256.OPT4 gp160 (accession #PZ137521) and CAP256.OPT4-S gp160 (accession #PZ137522) have been deposited in GenBank.

### HIV Env Protein Production

HIV-1 envelope was expressed and purified as previously described (61). Briefly, Freestyle 293F cells were diluted at the time of transfection to 1.25 × 106 cells/mL with fresh Freestyle293 media in 500 mL batches. Co-transfection was performed with plasmid DNA (650 µg SOSIP trimer plasmid and 150 µg furin expressing plasmid per 1L of culture volume) complexed with 293fectin in Opti-MEM I. After 6 days, cell cultures were harvested by centrifugation of the cells for 60 minutes at 4000 rpm in a Sorvall tabletop centrifuge. Supernatant was filtered through 0.8 µm filter and concentrated to approximately 100 mL with Vivaflow 200 cassettes (Sartorius) with a 30 kDa MWCO. Concentrated cell-free harvest was filtered again with a 0.8 µm cellulose filter system. The clarified concentrate was applied to a 10 mL affinity column containing PGT145 or 2G12 conjugated CnBr-activated Sepharose resin, using an AKTA Pure set to a flow rate of 2 mL per min. Following column was washed in PBS with 0.05% sodium azide, Env trimers were eluted using 3M MgCl_2_, and immediately diluted with 5 volumes of 10 mM tris, pH8. The eluate was filtered with a 0.2 µm filter bottle system and concentrated to 2 mL with a centricon-70 10 kDa MCWO. To biotinylate avi-tagged Envs, 25 µM of avi-tagged Env was dialyzed into Tris pH8 for 2-16 h. Dialyzed Env was biotinylated in a BirA biotin-protein ligase reaction (Avidity LLC) for 5 hours with mild agitation at 30 °C. Biotinylated envelope was concentrated to 2.4 mL prior to size-exclusion chromatography. Approximately, two milliliters of concentrated protein were purified on a HiLoad 16/60 Superose 6 pg column (Cytvia) in 500 mM NaCl buffer with 10 mM Tris, pH 8 to isolate trimeric protein. All chromatography steps were conducted on an AKTA Pure (Cytvia). The trimeric Env fractions were pooled, filtered with a 0.2 µm spin filter column, and snap frozen for long-term storage at −80 °C.

### Sortase A conjugation of HIV-1 envelope trimers to ferritin nanoparticles

Env trimer conjugate nanoparticle production was performed as described previously (61, 62). Sortase A-tagged CAP256.wk34.c80.OPT4 soluble trimers were expressed with amino acids LPSTGG encoded at its C terminus in Freestyle293 cells. Envelopes were purified by affinity chromatography with Env trimer-specific antibody PGT145 and Superose6 16/60 column chromatography as stated above. Helicobacter pylori ferritin subunits were expressed with a GGGGG repeat sequence encoded at their N terminus. Ferritin was expressed in Freestyle 293F cells by transiently transfecting 650 μg of expression plasmid with 293fectin. Five days post transfection, ferritin nanoparticles were purified by nickel chromatography. His tags were cleaved off of the ferritin by HRV-3C cleavage overnight. Envelopes with a C-terminal sortase A tag and ferritin particles with a sortase A N-terminal tag were buffer-exchanged into 50 mM Tris, 150 mM NaCl, 5 mM CaCl2, pH 7.5. 120 μM SOSIP gp140 was mixed with 120 μM ferritin subunits and incubated with 100 μM sortase A overnight at room temperature. After incubation, conjugated particles were isolated from free ferritin or free SOSIP gp140 using a Superose6 16/60 size-exclusion chromatography followed by sephacryl S-500 10/300 size-exclusion chromatography. The final protein was snap frozen and stored in 10 mM Tris, 500 mM NaCl pH8.0 at -80°C.

### Conjugation of Spytagged HIV-1 envelope trimers to mi3-SpyCatcher

Env trimer conjugation to mi3-SpyCatcher production was performed as described previously (63). Briefly, Spy Tagged BF1266 soluble Env trimers were incubated with mI3_SpyCatcher in 150mM NaCl, 25mM Tris pH8.0 at a ratio of 3:1 for 16 hours at room temperature. Conjugated Env nanoparticles were isolated from mI3_SpyCatcher or free SOSIP gp140 using Sephacryl S-500 10/300 size-exclusion chromatography. The final protein was snap frozen and stored in 10 mM Tris, 500 mM NaCl pH8.0 at -80°C. Overall shape and structure of the nanoparticle was characterized by size exclusion chromatography and NSEM.

### Generation of Fluorescently labeled Soluble Trimers for Flow Cytometry

Stabilized HIV Env trimers with C-terminal AviTags were expressed in Freestyle 293F cells (ThermoFisher) and were purified and biotinylated as previously described (61). Purified biotinylated Envs were conjugated to streptavidin-BV421 (Biolegend 405225), streptavidin-AF647 (Invitrogen S32357), or streptavidin-VB515 (MiltenyiBiotec 130-108-993) by four additions of a 1:1 molar ratio of fluorochrome-labeled streptavidin (final 4:1 ratio). Each addition was incubated with Env for 30 min at 4°C followed by the next addition of fluorochrome-labeled streptavidin. Finally, a ratio of 4:1 of soluble BIO-200 biotin (Avidity) was added to the reaction to quench any unoccupied sites on streptavidin. The quality of the fluorescently labeled soluble Env trimers was assessed by flow cytometry using a Ramos B cell line expressing CH01 RUA IgM. Ramos cells lacking surface immunoglobulin expression were used as a negative control cell line.

### B Cell Isolation

Fresh FNA cells from the inguinal lymph node (iLN) were stained and subjected to FACS. Briefly, cells from FNA were washed with PBS 1% BSA, and treated with 1mL of ACK Lysis for 2 min at RT, if blood contamination was observed. Cells were stained for 30 min at 4°C with a cocktail of antibodies in PBS 1% BSA against CD3 (clone SP34-2, BD 552852), IgD (polyclonal, Southern Biotech 2030-09), CD38 (clone OKT10, Caprico Biotechnologies1008206), CD21 (clone B-ly4, BD Biosciences 561374) IgG (Clone G18-145, BD Biosciences, 561297), CD71 (Clone L01.1, BD Biosciences, 746247) CD14 (clone M5E2, BioLegend 301832), IgM (G20-127, BD Biosciences, 562977), CD16 (Clone 3G8, BD Biosciences 563692), CD27 (Clone 0323, BioLegend 302834), CD20 (2H7, BioLegend 302356), BV421-, AF647, and/or VB515-conjugated soluble Env trimer. For cryopreserved peripheral blood mononuclear cells (PBMCs), 5-10 × 10^6^ cells were thawed and washed with RPMI 10% FBS. PBMCs were B cell enriched using EasySep Non-Human Primate B Cell Isolation Kit (Stem cell 100-0345) following manufacturer instructions. Enriched B cells were stained for 30 min with a cocktail of antibodies against CD3 (clone SP34-2, BD 552852), IgD (polyclonal, Southern Biotech 2030-09), CD38 (clone OKT10, Caprico Biotechnologies1008206), CD21 (clone B-ly4, BD Biosciences 561374), CD14 (clone M5E2, BioLegend 301832), CD16 (Clone 3G8, BD Biosciences 563692), CD27 (Clone 0323, BioLegend 302834), CD20 (2H7, BioLegend 302356), BV421-, AF647, and/or VB515-conjugated soluble Env trimer. Enriched B cells were washed with PBS 1%BSA, filtered through a sterile 100 μm strainer. Viability dye 7-amino-actinomycin D (7AA-D; ThermoFisher 00-6993-50) was added immediately prior to sorting cells on a BD FACS Symphony S6 Cell Sorter. Antigen-specific CD3-/CD14-/CD16-/CD20+/IgD-B cells were sorted into cell lysis buffer and 5X first-strand synthesis buffer in individual wells of a 96-well PCR plate. Plates were frozen on dry ice and ethanol immediately and stored at −80 °C until reverse transcription of RNA. OPT4-AF647+/VB515+ (OPT4++) specific B cells were sorted from FNA cells at weeks 15, 24, and 31 and PBMCs at week 12. OPT4-S-AF647+/VB515+ (OPT4-S++) Env-specific B cells were sorted from PBMC at week 53. BF1266-AF647+/BF1266-VB515+/BF1266.DKO-BV421-negative (BF1266++/BF1266DKO-) B cells were sorted from FNA cells at week 82 and 86 and PBMCs week 86.

### Rhesus Immunoglobulin RT-PCR

Immunoglobulin genes were amplified as previously described (64). Immunoglobulin genes from a single B cell were reverse transcribed with Superscript III (ThermoFisher) and constant region-specific reverse primers. Five microliters of complementary DNA were used for two rounds of nested PCR. PCR amplicons were purified and sequenced with 4 μM of forward and reverse primers. Contigs of the forward and reverse antibody sequences were assembled using the DHVI automated sequence analysis pipeline (DHVI ASAP).

### Enzymatically-Digested Antibody Fabs

Antibody Fabs were generated by papain digestion using the Pierce Fab Preparation Kit. The manufacturer’s protocol was followed except IgG antibodies were dialyzed into PBS for 2h prior to digestion and IgG was digested for 16–18 h. SDS-PAGE and Coomassie staining was used to examine the digestion of IgG into Fabs. Fabs were run through a Superdex200 10/300 column in 25mM Citric Acid with 125mM NaCl pH6 to remove any aggregates. Fabs were stored frozen at −80 °C in 25mM Citric Acid, 125mM NaCl pH 6 Buffer.

### Site-Directed Mutagenesis

OPT4-S, CAM13RRK, BF1266, and C.1080 Env-encoding plasmids were mutated to introduce V2 apex knockout substitutions N160K and R169E. CH01 RUA and PG16 RUA IgG plasmids were mutated to encode Fabs. Mutations were introduced using the QuikChange Lightning (Agilent) kit, following manufacturer recommended reaction conditions. Oligos introducing site-specific mutations were designed with the QuikChange Primer Design Tool (Agilent), synthesized and purified via standard desalting by Integrated DNA Technologies. Cloning was performed in XL-10 gold cells and sequence integrity was determined with Sanger sequencing (Azenta) and subsequent DNA alignment using Geneious (BioMatters).

### Recombinant Antibody and Fab Production

Monoclonal antibodies and Fabs were encoded in heavy and light chain plasmids and co-transfected into Expi293F (ThermoFisher) cells using expifectamine (ThermoFisher). Briefly, Expi293F cells were diluted to 2.5 × 10^6^ cells/mL with fresh Expi293 media in 100 mL batches and incubated for four hours prior to transfection. Fifty micrograms of each plasmid were mixed with OPTI-MEM I for 5 min and then added to expifectamine diluted in OPTI-MEM I. After 20 minutes, the DNA plus expifectamine was then added to the transfection culture. Expifectamine kit enhancers were added approximately 16–18 hours post-transfection and cultures were incubated for 5 days before harvest. The cultures were centrifuged at 4000 rpm in a Sorval tabletop centrifuge and the collected supernatant was filtered through 0.8 μm filter. The cell-free supernatant was incubated overnight at 4°C with either protein A resin for IgG1 or LambdaFabSelect or KappaSelect resin for lambda or kappa containing Fabs, respectively. The bead slurry was centrifuged at 1200 rpm for 10 minutes. The supernatant was aspirated from the pelleted resin. The resin was transferred to a gravity flow column, where the resin was washed with 20 mM Tris (pH 7) and 350 mM NaCl buffer. Fab or antibody was eluted with 2.5% acetic acid. The eluate was neutralized with Trizma (pH 8) and buffer exchanged into 25 mM citrate and 125 mM NaCl (pH 6) buffer through repetitive centrifugation in a Vivaspin Turbo-15 concentrator. Final protein was aliquoted and stored at −80 °C.

### IgG purification

2mL of plasma was heat-inactivated and centrifuged at 15,000 rpm for 5 min. Supernatant was collected and purified using Protein A/Protein G Sepharose Gravitrap kit (Cytiva 28985256), concentrated in 30 kDa centrifugal filters (Sigma UFC903024), and buffer exchanged into PBS. IgG concentration was determined with a Qubit Protein Assay kit (Invitrogen Q33212).

### Negative stain epitope mapping and data analysis

For monoclonal epitope mapping, IgG was mixed with OPT4-S Env trimer at a 1.5:1 ratio, diluted to 0.4 mg/ml nominal trimer concentration by adding buffer (150 mM Na2SO4, 2.5 mM NaCl, 2.8 NaN3, 20 mM HEPES, pH 7.4), and incubated 1 h at room temperature. For polyclonal epitope mapping (EMPEM), 1 mg of polyclonal Fab was mixed with 20 µg of OPT4-S, incubated overnight at 4 °C, and fractionated over a Superose 6 Increase 10/300 column the following day. Fractions corresponding to the presumed complex peak were pooled and concentrated in a 100-kDa MWCO spin concentrator. Subsequent processing was similar for polyclonal and monoclonal samples. A portion of the sample was diluted 1:1 in buffer containing 16 mM glutaraldehyde, fixed 5 min, quenched by adding 1 M Tris (pH 7.4) to give 80 mM final Tris concentration, and incubated for 5 min. Quenched sample was then diluted 1:1 with buffer and negatively stained by applying 7 µl of sample to a glow-discharged carbon-coated EM grid for 10-12 second, blotting, staining with 2 g/dL uranyl formate for 1 min, then blotting and air-drying. Grids were examined on a Philips EM420 electron microscope operating at 120 kV and nominal magnification of 49,000x, and ∼100 or ∼300 images (monoclonal vs polyclonal) were collected on a 76 Mpix CCD camera at 2.4 Å/pixel. Images were analyzed by automatic particle picking, 2D and 3D class averages; and 3D reconstructions were calculated using standard protocols with Relion 3.0 (65). For polyclonal samples, 3D classifications were used to estimate the relative occupancy at each epitope, by tallying the number of particles in each 3D class.

### Fab-Env complex formation, negative staining, and data analysis

For monoclonal Fabs, 36 μg of Fab was mixed with 10 μg OPT4-S Env trimer in a total volume of 100 μL of HEPES-buffered saline (HBS) and incubated overnight at 4 °C. Fab-trimer complexes were then diluted with 400 μL of HBS with 10 mM glutaraldehyde, incubated for 5 minutes at room temperature, and quenched by addition of 1 M Tris to 80 mM final concentration. Quenched samples were then concentrated with a 100-kDA molecular weight cutoff spin concentrator, which allows excess unbound Fabs to pass the filter and retains the Fab-trimer complex. Concentrated sample was then diluted and negatively stained as described above. Negatively stained grids were examined on a Philips EM420 electron microscope operating at 120 kV, 49,000x nominal magnification, and ∼0.5 μm defocus. Images were acquired on a 76-megapixel CCD camera, corresponding to a nominal calibration of 2.4 Å/pixel. Datasets were typically ∼100 images for monoclonal samples. Image analysis was performed with standard protocols in Relion65, beginning with automated particle picking, followed with two rounds of 2D classification/selection, and then by 1–2 rounds of 3D classification/selection to discard junk particles and select Fab-bound trimer particles. Monoclonal samples, particle stacks from well-resolved 3D classes were chosen and final 3D refinements with post-processing performed.

### Biolayer interferometry binding kinetic assays

Biolayer interferometry (BLI) was performed on an Octet Red96e system (Sartorius) at 30°C with an orbital shake speed of 1000 rpm. Assays were performed in flat bottom 96-well assays plates (Greiner) using 0.22µm-filtered Phosphate buffered saline supplemented with 0.05% Tween 20 and 0.1% bovine serum albumin (PBS-T-BSA). Binding kinetics experiment were performed with Env immobilized to hydrated streptavidin sensor tips at concentrations exhibiting a linear loading phase over 300 s. C.1080 and BF1266 Envs were tested at 1.25 μg/mL. OPT4-S Env was tested at 2.5 μg/mL. Env was submerged into two-fold serial dilutions of Fabs, starting at a Fab concentration of 1000μM for C1080 and BF1266 Envs and 100 μM for OPT4-S Env. Association occurred for 300 seconds followed by a 600-second dissociation step in PBS-T-BSA. The binding response sensorgram curves were globally fitted using a 1:1 binding model and the rate constants k_a_, k_d_ and K_d_ were calculated from curves for four or more Fab concentrations using Data Analysis HT 12.0 software (Forte Bio).

### Indirect ELISA Using Env Trimers

Corning 384-well plates were coated overnight at 4°C with 15 µL of streptavidin diluted to 2 µg/mL in 0.1M NaCO3. Plates were washed with PBS-T (PBS +0.05% Tween-20) and wells blocked for 1 hour at room temperature with 40 µL blocking buffer (SuperBlock consisting of PBS, 15% goat serum, 4% whey protein, 0.05% Tween-20). After blocking, plates were washed, then incubated with 10 µL per well of biotinylated Env diluted to 2 µg/mL in blocking buffer for 1 hour. Beginning at a concentration of 50 µg/mL monoclonal antibodies were serially diluted in SuperBlock. Similarly, serum was diluted 1:30 in SuperBlock and serially diluted. Plates were washed and incubated with 10 µL of a serial dilution of monoclonal antibody or rhesus macaque serum for 1.5 hours. Wells were washed two times with PBS-T, then incubated for 1 hour with 10μL of 1:12,000 dilution of goat anti-human IgG-HRP (Jackson ImmunoResearch Cat. No:109-035-098) or 1:30,000 dilution of mouse anti-rhesus IgG-HRP (Southern Biotech Cat. No: 4700–05) in SuperBlock lacking sodium azide. Plates were washed two times, then developed with 20 µL per well of tetramethylbenzidine peroxidase substrate for 15 minutes, then quenched with equal volume 1% HCl. Absorbance was measured at 450 nm on a SpectraMax 340 PC. Binding titers were analyzed as area-under-curve of the log-transformed concentration curve.

### Virus neutralization assays

Virus neutralization assays were performed with HIV-1 Env-pseudotyped viruses as described previously (66). SHIV and Env-pseudotyped virus stocks were generated by transfection of HEK 293T/17 cells, as previously described (67). Neutralizing antibody activity was measured in 96-well culture plates using Tat-regulated luciferase (Luc) reporter gene expression to quantify inhibition of virus infection in TZM-bl cells. TZM-bl cells were obtained from the NIH AIDS Research and Reference Reagent Program, as contributed by John Kappes and Xiaoyun Wu. Serum samples were heat-inactivated at 56 °C for 30 minutes prior to assay. Test serum samples were diluted over a range of 1:20 to 1:43740 in cell culture medium and pre-incubated with virus (about 150,000 relative light unit equivalents) for 1 hour at 37 °C before addition of cells. Similarly, mAbs were serially diluted three-fold in cell culture medium starting at 300µg/mL and pre-incubated with virus under the above conditions. Following a 48-hour incubation, cells were lysed and Luc activity determined using a microtiter plate luminometer and BriteLite Plus Reagent (Perkin Elmer). Neutralization titers are the sample dilution (for serum) or antibody concentration (for mAbs) at which relative luminescence units (RLU) were reduced by 50% (ID_50_) compared to RLU in virus only control wells.

### Bulk NGS processing and analysis of V2-apex specific clonal lineages

The R1 and R2 FASTQ files for each rhesus macaque (RM) were first polyG trimmed using fastp (https://github.com/opengene/fastp) and then merged with the flash program. Merged reads were quality filtered such that retained reads contained at least 95% of bases with a Phred quality score ≥30. Retained reads were converted to FASTA format and reverse complemented for downstream analysis.

Confident immunoglobulin heavy-chain sequences were identified using a rhesus macaque–specific chain classifier (68). Sequences sharing the same UMI were collapsed into a consensus sequence using a custom Python script. Prior to consensus building, sequences with lengths outside the mean ± one standard deviation (calculated per UMI) were discarded. For each position, the majority base was called if present at >50%; otherwise, an N was assigned. Consensus sequences containing more than 10 Ns were removed. UMIs represented by a single sequence were retained without collapsing.

UMI-collapsed sequences were assigned to isotypes by aligning sequence regions 3′ of the VDJ junction to a reference set of *Macaca mulatta* constant regions (68). VDJ boundaries were initially identified using the Cloanalyst (69) software package.

IgD and IgM transcripts were used to infer per-monkey immunoglobulin rearrangement and mutation parameters using partis (70, 71) and the Macaca mulatta OGRDB germline sequences (release 2026-02-05; Zenodo: 10.5281/zenodo.18172063) (72). These learned parameters were then applied to annotate and partition the combined set of the IgG NGS sequences, the sorted 10X sequences (wk 12), non-sorted 10X sequences (wk 53), and the sorted single-cell PCR sequences.

### Clonal lineage UCA and paired tree inference

All sorted single-cell PCR sequences from NHP 192254 were annotated and partitioned into clonal lineages using partis (70, 71). Parameter estimation was performed using a directory built from these sorted single-cell PCR sequences combined with 10X genomic repertoire sequences from the same PTID collected at week 235, using Macaca mulatta OGRDB germline sequences (release 2026-02-05; Zenodo: 10.5281/zenodo.18172063) (72). All sequences in the heavy chain clone that included the experimentally validated V2-apex targeting mAbs DH2050 lineage were selected for UCA inference and paired lineage reconstruction. The corresponding paired kappa light chain sequences were verified to share the same V gene, J gene, and CDR3 length, supporting correct pairing and clonal assignment. Unmutated common ancestor (UCA) inference was performed independently for heavy and kappa chains using linearham (73), based on the inferred clonal members and using the same parameter directory as for initial annotation and partitioning. Inference was run with parameters --mcmc-iter=10000, --mcmc-thin=10, and --tune-iter=1. A single kappa-chain UCA was inferred. For the heavy chain, multiple putative

UCA sequences were returned; the UCA with the highest posterior probability was selected for downstream analyses. Paired phylogenetic trees were inferred using dowser (tree-building method: RAxML), with the inferred heavy and kappa UCAs specified as the germline sequences. The heavy chain tip sequences and UCA were jointly aligned to account for the presence of insertions and deletions prior to tree inference (74).

### Cryo-electron microscopy

HIV-1 envelope at a final concentration of 1.22 mg/mL was mixed with VRC34.01 fab and vaccine-elicited antibody at a 1:4:3 molar ratio in PBS buffer and incubated at room temperature for 30 minutes. The VRC34.01 antibody was added to prevent the preferred orientation observed on vitrified Cryo-EM grids when the envelope was incubated only with the vaccine-elicited antibodies. The Quantifoil R1.2/1.3 Au 300 grids (Electron Microscopy Sciences) were glow-discharged at 15 mA for 15 seconds using PELCO easiGlow™ from TED PELLA, INC. Vitrification was carried out using the Leica EM GP2 (Leica Microsystems), maintaining 95% Humidity and 22 °C in the chamber. A 3 µL volume of the sample was applied to the glow-discharged grids and incubated for 30 seconds before blotting for 1.5 seconds. Then the grids were plunged immediately into liquid ethane and vitrified in liquid nitrogen. Micrographs were collected using a 100 keV TFS Tundra electron microscope with a Falcon C detector at 180kX magnification (1.47-pixel size). Those micrographs were processed using CryoSPARC (75) to generate the Cryo-EM maps. Patch motion correction and Patch CTF estimation were performed on micrographs, followed by Blob picker, extraction of micrographs in 256 px box size, 2D classification, and 3D construction. Non-uniform refinement and local refinements were performed to further refine the maps. Models of the envelope and Fabs were fitted to the maps and refined further using ChimeraX (PMID: (76), Phenix (77) and Coot (78) software.

## REFERENCES

1. C. Copeland et al., The HIV Epidemic in the United States -Epidemiological Projections and Public Economic Impact of Achieving Zero Transmission Goals. Clinicoecon Outcomes Res 17, 755–769 (2025).

2. L. Corey et al., Two Randomized Trials of Neutralizing Antibodies to Prevent HIV-1 Acquisition. N Engl J Med 384, 1003–1014 (2021).

3. A. Pegu et al., A Meta-analysis of Passive Immunization Studies Shows that Serum-Neutralizing Antibody Titer Associates with Protection against SHIV Challenge. Cell Host Microbe 26, 336–346 e333 (2019).

4. L. M. Walker et al., Broad and potent neutralizing antibodies from an African donor reveal a new HIV-1 vaccine target. Science 326, 285–289 (2009).

5. X. Wu et al., Rational design of envelope identifies broadly neutralizing human monoclonal antibodies to HIV-1. Science 329, 856–861 (2010).

6. B. F. Haynes et al., Strategies for HIV-1 vaccines that induce broadly neutralizing antibodies. Nat Rev Immunol 23, 142–158 (2023).

7. P. D. Kwong, J. R. Mascola, HIV-1 Vaccines Based on Antibody Identification, B Cell Ontogeny, and Epitope Structure. Immunity 48, 855–871 (2018).

8. E. Parker Miller, M. T. Finkelstein, M. C. Erdman, P. C. Seth, D. Fera, A Structural Update of Neutralizing Epitopes on the HIV Envelope, a Moving Target. Viruses 13, (2021).

9. J. Gorman et al., Structures of HIV-1 Env V1V2 with broadly neutralizing antibodies reveal commonalities that enable vaccine design. Nat Struct Mol Biol 23, 81–90 (2016).

10. J. H. Lee et al., A Broadly Neutralizing Antibody Targets the Dynamic HIV Envelope Trimer Apex via a Long, Rigidified, and Anionic beta-Hairpin Structure. Immunity 46, 690–702 (2017).

11. R. D. Mason et al., Structural development of the HIV-1 apex-directed PGT145-PGDM1400 antibody lineage. Cell Rep 44, 115223 (2025).

12. J. S. McLellan et al., Structure of HIV-1 gp120 V1/V2 domain with broadly neutralizing antibody PG9. Nature 480, 336–343 (2011).

13. N. A. Doria-Rose et al., New Member of the V1V2-Directed CAP256-VRC26 Lineage That Shows Increased Breadth and Exceptional Potency. J Virol 90, 76–91 (2016).

14. D. Sok et al., Recombinant HIV envelope trimer selects for quaternary-dependent antibodies targeting the trimer apex. Proc Natl Acad Sci U S A 111, 17624–17629 (2014).

15. M. Bonsignori et al., Analysis of a clonal lineage of HIV-1 envelope V2/V3 conformational epitope-specific broadly neutralizing antibodies and their inferred unmutated common ancestors. J Virol 85, 9998–10009 (2011).

16. N. A. Doria-Rose et al., Developmental pathway for potent V1V2-directed HIV-neutralizing antibodies. Nature 509, 55–62 (2014).

17. E. Landais et al., HIV Envelope Glycoform Heterogeneity and Localized Diversity Govern the Initiation and Maturation of a V2 Apex Broadly Neutralizing Antibody Lineage. Immunity 47, 990–1003 e1009 (2017).

18. E. M. Cale et al., Virus-like Particles Identify an HIV V1V2 Apex-Binding Neutralizing Antibody that Lacks a Protruding Loop. Immunity 46, 777–791 e710 (2017).

19. B. Julg et al., Broadly neutralizing antibodies targeting the HIV-1 envelope V2 apex confer protection against a clade C SHIV challenge. Sci Transl Med 9, (2017).

20. A. Pegu et al., Neutralizing antibodies to HIV-1 envelope protect more effectively in vivo than those to the CD4 receptor. Sci Transl Med 6, 243ra288 (2014).

21. E. Landais et al., Broadly Neutralizing Antibody Responses in a Large Longitudinal Sub-Saharan HIV Primary Infection Cohort. PLoS Pathog 12, e1005369 (2016).

22. L. M. Walker et al., A limited number of antibody specificities mediate broad and potent serum neutralization in selected HIV-1 infected individuals. PLoS Pathog 6, e1001028 (2010).

23. W. B. Williams et al., Vaccine induction of heterologous HIV-1-neutralizing antibody B cell lineages in humans. Cell 187, 2919–2934 e2920 (2024).

24. B. F. Haynes, D. R. Burton, J. R. Mascola, Multiple roles for HIV broadly neutralizing antibodies. Sci Transl Med 11, (2019).

25. B. F. Haynes, G. Kelsoe, S. C. Harrison, T. B. Kepler, B-cell-lineage immunogen design in vaccine development with HIV-1 as a case study. Nat Biotechnol 30, 423–433 (2012).

26. B. S. Briney, J. R. Willis, J. E. Crowe, Jr., Human peripheral blood antibodies with long HCDR3s are established primarily at original recombination using a limited subset of germline genes. PLoS One 7, e36750 (2012).

27. J. R. Willis et al., Human immunoglobulin repertoire analysis guides design of vaccine priming immunogens targeting HIV V2-apex broadly neutralizing antibody precursors. Immunity 55, 2149–2167 e2149 (2022).

28. P. L. Moore, J. Gorman, N. A. Doria-Rose, L. Morris, Ontogeny-based immunogens for the induction of V2-directed HIV broadly neutralizing antibodies. Immunol Rev 275, 217–229 (2017).

29. R. Andrabi et al., Identification of Common Features in Prototype Broadly Neutralizing Antibodies to HIV Envelope V2 Apex to Facilitate Vaccine Design. Immunity 43, 959–973 (2015).

30. F. Bibollet-Ruche et al., A Germline-Targeting Chimpanzee SIV Envelope Glycoprotein Elicits a New Class of V2-Apex Directed Cross-Neutralizing Antibodies. mBio 14, e0337022 (2023).

31. R. Andrabi et al., The Chimpanzee SIV Envelope Trimer: Structure and Deployment as an HIV Vaccine Template. Cell Rep 27, 2426–2441 e2426 (2019).

32. M. Medina-Ramirez et al., Design and crystal structure of a native-like HIV-1 envelope trimer that engages multiple broadly neutralizing antibody precursors in vivo. J Exp Med 214, 2573–2590 (2017).

33. A. R. Ghosh et al., Rapidly acquired HIV-1 neutralization breadth in a rhesus V2 apex knockin mouse model after a single bolus immunization. Sci Immunol 11, eadz5064 (2026).

34. E. Melzi et al., Membrane-bound mRNA immunogens lower the threshold to activate HIV Env V2 apex-directed broadly neutralizing B cell precursors in humanized mice. Immunity 55, 2168–2186 e2166 (2022).

35. J. E. Voss et al., Elicitation of Neutralizing Antibodies Targeting the V2 Apex of the HIV Envelope Trimer in a Wild-Type Animal Model. Cell Rep 21, 222–235 (2017).

36. H. Duan et al., Vaccine elicitation of HIV-1 neutralizing antibodies against both V2 apex and fusion peptide in rhesus macaques. Cell Rep 45, 116905 (2026).

37. J. Guenaga et al., Vaccination generates broadly cross-neutralizing antibodies to the HIV Env apex. Nature, (2026).

38. K. M. Ma et al., HIV broadly neutralizing antibody precursors to the Apex epitope induced in nonhuman primates. Sci Immunol 10, eadt6660 (2025).

39. N. Mishra et al., Germline-targeting HIV immunogen induces cross-neutralizing antibodies in outbred macaques. Immunity 59, 1140–1160 e1111 (2026).

40. K. O. Saunders et al., Vaccine Induction of Heterologous Tier 2 HIV-1 Neutralizing Antibodies in Animal Models. Cell Rep 21, 3681–3690 (2017).

41. L. Marchitto et al., Enhanced Priming Leads to Rapid Induction of Broadly Neutralizing HIV-1 Apex Antibodies. Nature, (2026).

42. R. Habib et al., Env-antibody coevolution identifies B cell priming as the principal bottleneck to HIV V2 apex broadly neutralizing antibody development. Sci Immunol 11, eadz3933 (2026).

43. R. S. Roark et al., Structural and genetic basis of HIV-1 envelope V2 apex recognition by rhesus broadly neutralizing antibodies. J Exp Med 222, (2025).

44. R. Pejchal et al., Structure and function of broadly reactive antibody PG16 reveal an H3 subdomain that mediates potent neutralization of HIV-1. Proc Natl Acad Sci U S A 107, 11483–11488 (2010).

45. M. Pancera et al., Structural basis for diverse N-glycan recognition by HIV-1-neutralizing V1-V2-directed antibody PG16. Nat Struct Mol Biol 20, 804–813 (2013).

46. R. S. Roark et al., Recapitulation of HIV-1 Env-antibody coevolution in macaques leading to neutralization breadth. Science 371, (2021).

47. T. Hagglof et al., Continuous germinal center invasion contributes to the diversity of the immune response. Cell 186, 147–161 e115 (2023).

48. L. P. Deimel et al., Clonal expansion and diversification of germinal center and memory B cell responses to booster immunization in primates. Cell Rep 44, 116142 (2025).

49. K. M. Cirelli et al., Slow Delivery Immunization Enhances HIV Neutralizing Antibody and Germinal Center Responses via Modulation of Immunodominance. Cell 177, 1153–1171 e1128 (2019).

50. R. Habib et al., Deep mining of the human antibody repertoire identifies frequent and genetically diverse CDRH3 topologies targetable by vaccination. Proc Natl Acad Sci U S A 123, e2532810123 (2026).

51. J. Gorman et al., Structure of Super-Potent Antibody CAP256-VRC26.25 in Complex with HIV-1 Envelope Reveals a Combined Mode of Trimer-Apex Recognition. Cell Rep 31, 107488 (2020).

52. H. N. Eisen, G. W. Siskind, Variations in Affinities of Antibodies during the Immune Response. Biochemistry 3, 996–1008 (1964).

53. D. Paus et al., Antigen recognition strength regulates the choice between extrafollicular plasma cell and germinal center B cell differentiation. J Exp Med 203, 1081–1091 (2006).

54. L. Mesin et al., Restricted Clonality and Limited Germinal Center Reentry Characterize Memory B Cell Reactivation by Boosting. Cell 180, 92–106 e111 (2020).

55. A. Schiepers, M. F. L. Van’t Wout, A. Hobbs, L. Mesin, G. D. Victora, Opposing effects of pre-existing antibody and memory T cell help on the dynamics of recall germinal centers. Immunity 57, 1618–1628 e1614 (2024).

56. Y. Zhang et al., Germinal center B cells govern their own fate via antibody feedback. J Exp Med 210, 457–464 (2013).

57. J. H. Lee et al., Long-primed germinal centres with enduring affinity maturation and clonal migration. Nature 609, 998–1004 (2022).

58. K. Wagh et al., Signature based vaccines induce V2 apex broadly neutralizing antibodies in preparation, (2026).

59. E. T. Crooks et al., Engineering well-expressed, V2-immunofocusing HIV-1 envelope glycoprotein membrane trimers for use in heterologous prime-boost vaccine regimens. PLoS Pathog 17, e1009807 (2021).

60. C. A. Bricault et al., HIV-1 Neutralizing Antibody Signatures and Application to Epitope-Targeted Vaccine Design. Cell Host Microbe 25, 59–72 e58 (2019).

61. K. O. Saunders et al., Targeted selection of HIV-specific antibody mutations by engineering B cell maturation. Science 366, (2019).

62. K. O. Saunders et al., Vaccine induction of CD4-mimicking HIV-1 broadly neutralizing antibody precursors in macaques. Cell 187, 79–94 e24 (2024).

63. T. U. J. Bruun, A. C. Andersson, S. J. Draper, M. Howarth, Engineering a Rugged Nanoscaffold To Enhance Plug-and-Display Vaccination. ACS Nano 12, 8855–8866 (2018).

64. K. O. Saunders et al., Stabilized HIV-1 envelope immunization induces neutralizing antibodies to the CD4bs and protects macaques against mucosal infection. Sci Transl Med 14, eabo5598 (2022).

65. J. Zivanov et al., New tools for automated high-resolution cryo-EM structure determination in RELION-3. Elife 7, (2018).

66. M. Li et al., Human immunodeficiency virus type 1 env clones from acute and early subtype B infections for standardized assessments of vaccine-elicited neutralizing antibodies. J Virol 79, 10108–10125 (2005).

67. H. Li et al., Envelope residue 375 substitutions in simian-human immunodeficiency viruses enhance CD4 binding and replication in rhesus macaques. Proc Natl Acad Sci U S A 113, E3413–3422 (2016).

68. S. Holmes et al., Neonatal immunity associated with heterologous HIV-1 neutralizing antibody induction in SHIV-infected Rhesus Macaques. Nat Commun 15, 10302 (2024).

69. T. B. Kepler, Reconstructing a B-cell clonal lineage. I. Statistical inference of unobserved ancestors. F1000Res 2, 103 (2013).

70. D. K. Ralph, F. A. t. Matsen, Consistency of VDJ Rearrangement and Substitution Parameters Enables Accurate B Cell Receptor Sequence Annotation. PLoS Comput Biol 12, e1004409 (2016).

71. D. K. Ralph, F. A. t. Matsen, Per-sample immunoglobulin germline inference from B cell receptor deep sequencing data. PLoS Comput Biol 15, e1007133 (2019).

72. A. Peres et al., Population-level genomic analysis of immunoglobulin loci variation in rhesus macaques reveals extensive germline diversity. Immunity 59, 213–228 e216 (2026).

73. A. Dhar, D. K. Ralph, V. N. Minin, F. A. t. Matsen, A Bayesian phylogenetic hidden Markov model for B cell receptor sequence analysis. PLoS Comput Biol 16, e1008030 (2020).

74. C. G. Jensen, J. A. Sumner, S. H. Kleinstein, K. B. Hoehn, Inferring B Cell Phylogenies from Paired H and L Chain BCR Sequences with Dowser. J Immunol 212, 1579–1588 (2024).

75. A. Punjani, J. L. Rubinstein, D. J. Fleet, M. A. Brubaker, cryoSPARC: algorithms for rapid unsupervised cryo-EM structure determination. Nat Methods 14, 290–296 (2017).

76. E. C. Meng et al., UCSF ChimeraX: Tools for structure building and analysis. Protein Sci 32, e4792 (2023).

77. D. Liebschner et al., Macromolecular structure determination using X-rays, neutrons and electrons: recent developments in Phenix. Acta Crystallogr D Struct Biol 75, 861–877 (2019).

78. P. Emsley, B. Lohkamp, W. G. Scott, K. Cowtan, Features and development of Coot. Acta Crystallogr D Biol Crystallogr 66, 486–501 (2010).

