## Supplemental Figures for "Rapid vaccine induction of macaque HIV-1 V2 Apex broadly neutralizing antibodies with immunogenetic signatures that are potentially translatable to humans"

**Supplemental Figure 1**

**A.** OPT4-scNP

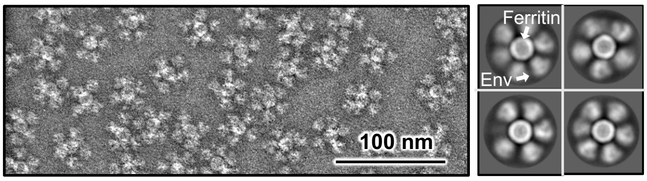


Box =480 Å

**B.** BF1266-scNP

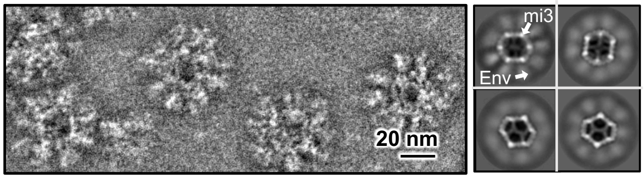


Box =480 Å

**C.**

| Envelopes: | CH065 | 2G12 | PGT145 | VRC26.25 | CH01 | PG16 | PG9 | PCT64 | PG9_RUA | PG16_RUA | CH01_RUA | VRC26.25_RUA | 41328_UCA | RHA1_UCA | 40591_UCA | 42056.P1_UCA | V031_UCA | V033_UCA |
|---|---|---|---|---|---|---|---|---|---|---|---|---|---|---|---|---|---|---|
| | | | V2-apex bnabs | | | | | | V2-apex bnabs precursors | | | | | | | | | |
| OPT4 | 0 | 11.5 | 13.5 | 13.0 | 14.4 | 15.8 | 14.1 | 15.0 | 8.2 | 10.2 | 13.5 | 0 | 10.8 | 7.5 | 4.8 | 0.1 | 0.8 | 0 |
| OPT4.DKO | 0 | 12.0 | 0.2 | 11.8 | 0 | 0.2 | 0.9 | 0.1 | 0 | 0 | 0 | 0 | 0 | 0 | 0 | 0 | 0 | 0 |
| OPT4-S | 0 | 10.9 | 13.5 | 15.1 | 13.9 | 16.0 | 14.0 | 14.4 | 7.5 | 9.7 | 13.4 | 0 | 9.2 | 3.7 | 1.2 | 0 | 0 | 0 |
| OPT4-S.DKO | 0 | 10.0 | 0 | 12.0 | 0 | 0 | 0 | 0 | 0 | 0 | 0 | 0 | 0 | 0 | 0 | 0 | 0 | 0 |
| CAM13RRK | 0 | 12.0 | 15.2 | 14.9 | 14.0 | 15.8 | 13.6 | 14.9 | 12.3 | 13.5 | 9.8 | 0 | 7.1 | 0 | 0 | 0 | 0 | 0 |
| CAM13RRK.DKO | 0 | 8.9 | 0 | 0 | 0 | 0 | 0 | 0 | 0 | 0 | 0 | 0 | 0 | 0 | 0 | 0 | 0 | 0 |
| C.1080 | 0 | 7.9 | 12.6 | 13.4 | 8.7 | 14.3 | 12.2 | 9.8 | 6.8 | 0 | 0 | 0 | 0 | 0 | 0 | 0 | 0 | 2.2 |
| C.1080.DKO | 0 | 10.3 | 0 | 0.9 | 0 | 0.1 | 0 | 0 | 0 | 0 | 0 | 0 | 0 | 0 | 0 | 0 | 0 | 0 |
| BF1266 | 0 | 9.4 | 12.5 | 12.0 | 12.2 | 16.0 | 12.6 | 8.5 | 0.4 | 0.2 | 0.5 | 0 | 0 | 0 | 0 | 0 | 0 | 0 |
| BF1266.DKO | 0 | 10.5 | 0 | 0 | 0 | 0 | 0 | 0 | 0 | 0 | 0 | 0 | 0 | 0 | 0 | 0 | 0 | 0 |

Binding (log AUC): 0, 5, 10, 15

**D.**

| HIV-1 Env | C-strand motif |
|---|---|
| OPT4 | RDKRRK |
| OPT4_R189T | RDKRRK |
| C.1080 | RDKKQK |
| BF1226 | RDKKKK |
| Q23 | RDKRQK |
| T250 | RDKKKK |
| ZM233 | RDKKRK |
| 16055 | RDKKQK |
| 246F3 | RDKRQK |
| CM244 | RDKKQK |
| CE1176 | KDKKKN |
| CE0217 | KDKKKK |
| CAM13RRK | RDKRRK |
| SHIV.BG505 | RDKKQK |
| C4118 | RDRKQK |
| 1394C9 | RDKKKK |
| V703_2117 | RDKRRK |
| V703_2018 | RDKKRK |
| V703_2149 | KDRRRK |
| CE0393 | KDKKKK |
| 0921.v2.c14 | KDKKRK |
| 9004SS_A3_4 | RDKKQK |
| DU123_06 | RDKKQK |
| Q168.a2 | RDKRQK |
| 1394_C9G1 | RDKKKK |

**E.**

| Position | | | | | | |
|---|---|---|---|---|---|---|
| 166 | aa | R | K | G | T | others |
| | % | 68 | 15.2 | 4.5 | 2.6 | 9.7 |
| 167 | aa | D | N | G | others | |
| | % | 86.5 | 6.4 | 2.4 | 4.7 | |
| 168 | aa | K | R | others | | |
| | % | 86 | 11.5 | 2.5 | | |
| 169 | aa | K | V | R | Q | others |
| | % | 42.4 | 17.1 | 9 | 8.2 | 23.3 |
| 170 | aa | Q | K | R | E | others |
| | % | 50.7 | 31.4 | 9.0 | 2.9 | 6 |
| 171 | aa | K | Q | R | T | others |
| | % | 65.8 | 11.2 | 7.7 | 5.1 | 10.2 |

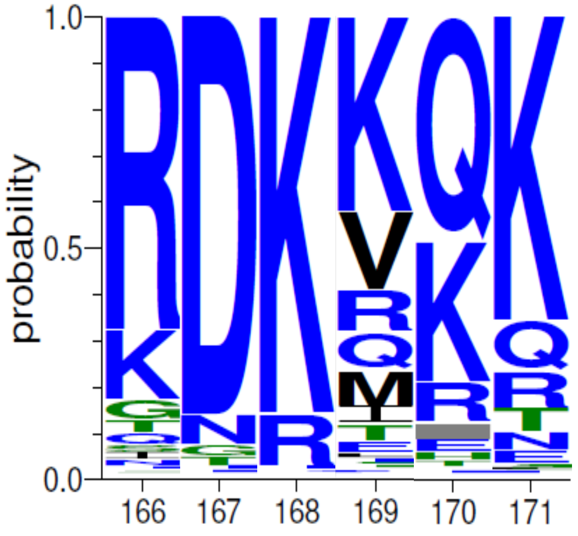

**Supplemental Figure 1. Nanoparticles imaging and soluble envelopes binding profile and structural characteristics related to Figure 1.**

(A) OPT4-scNP: Raw electron micrograph (left) and 2D class average (right). (B) BF1266-mi3NP: Raw electron micrograph (left) and 2D class average (right). (C) V2-apex bnabs and inferred precursors binding by ELISA against a panel of soluble envelopes trimers and their respective V2-apex double knock out (DKO – K169E, N160K) mutant. (D) Details of HIV-1 envelopes C-strand motif. (E) Table with the amino acid variation in HIV-1 envelopes at position 166 to 171 (C-strand motif).

**Supplemental Figure 2**

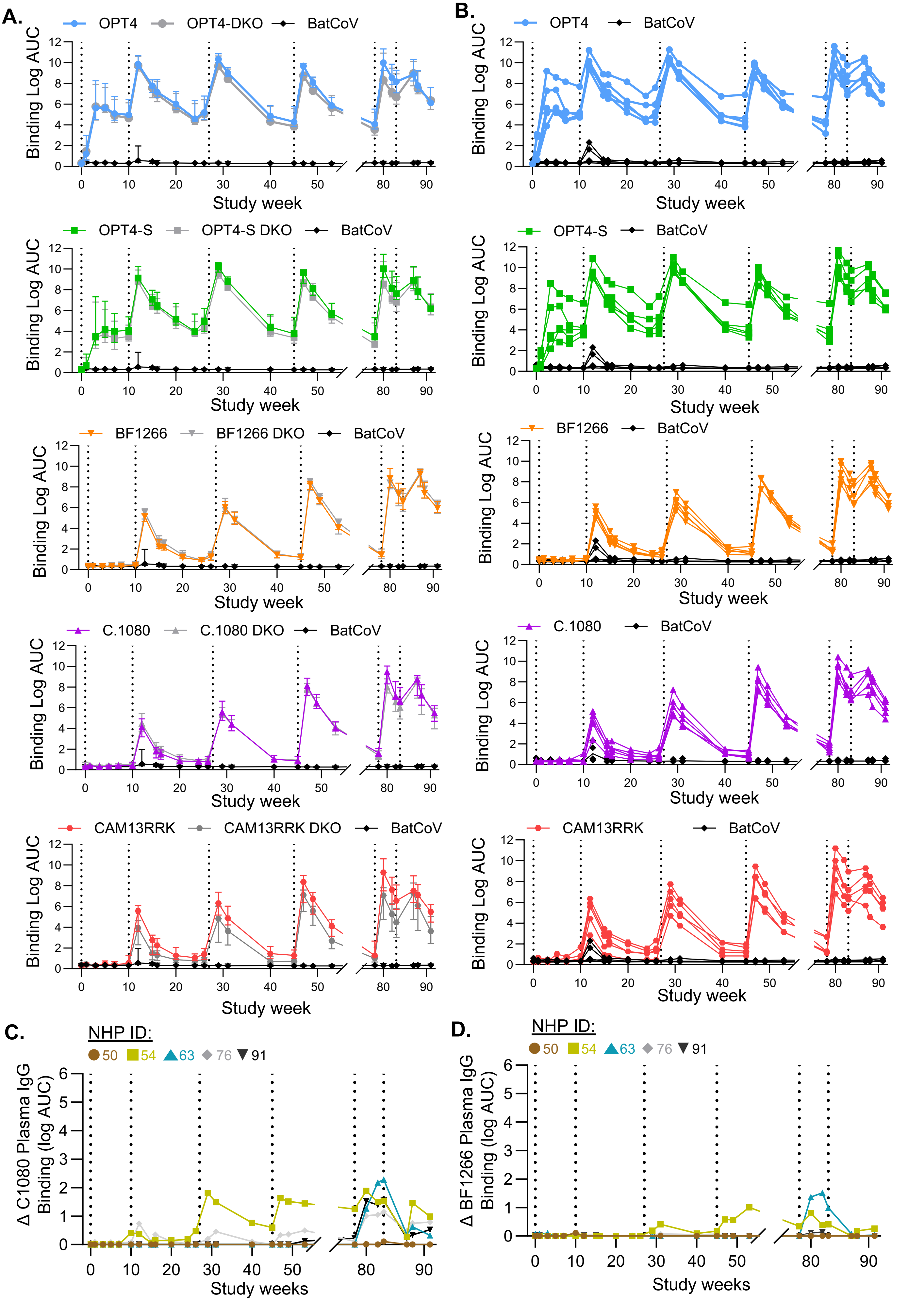

**Supplemental Figure 2. Rhesus macaques plasma binding profile related to Figure 1.**

(A) Rhesus macaques Plasma binding by ELISA against a panel of soluble envelopes trimers and their respective V2-apex double knock out (DKO – K169E, N160K) mutant. Results shown as Median log AUC and Interquartile range (B) Individual Rhesus macaques Plasma binding by ELISA against a panel of soluble envelopes trimers. (C) Differential plasma binding between C.1080 and C.1080 DKO soluble trimers. Results showed as median log AUC. (D) Differential plasma binding between BF1266 and BF1266 DKO soluble trimers. Results showed as median log AUC.

**A**

| Group | Subtype | Tier | Immunogen | OPT4-scNP | | | | | OPT4-scNP | | | | | OPT4-S mRNA | | | | |
|---|---|---|---|---|---|---|---|---|---|---|---|---|---|---|---|---|---|---|
| | | | NHP ID | **50** | **54** | **63** | **76** | **91** | **50** | **54** | **63** | **76** | **91** | **50** | **54** | **63** | **76** | **91** |
| | | | Study Week | **wk3** | | | | | **wk12** | | | | | **wk29** | | | | |
| | Subtype | Tier | MLV | <20 | <20 | <20 | <20 | <20 | <20 | <20 | <20 | <20 | <20 | <20 | <20 | <20 | <20 | <20 |
| Autologous Viruses | | | OPT4 | 164 | 299 | 1637 | 425 | 99 | 238949 | 98232 | 172592 | 199005 | 119303 | 334896 | 358166 | 366838 | 361141 | 371058 |
| | | | OPT4-S | <20 | <20 | <20 | <20 | <20 | 1280 | 5123 | 2976 | 912 | 1753 | 14017 | 45767 | 45208 | 35249 | 63291 |
| | | | OPT4-S.N160K | | | | | | 724 | 347 | 942 | 589 | 912 | 5176 | 8410 | 41288 | 36563 | 45249 |
| | | | OPT4-S.R169E | | | | | | 332 | 192 | 982 | 165 | 373 | 2005 | 5896 | 15333 | 15959 | 32938 |
| Viruses lacking N130 | A | 1B | Q23.17 | <20 | <20 | <20 | <20 | <20 | <20 | 422 | <20 | <20 | <20 | <20 | 33738 | <20 | <20 | <20 |
| | 02_AG | 2 | T250 | | | | | | 122 | <20 | 20 | <20 | <20 | <20 | 30 | <20 | <20 | <20 |
| | C | 2 | ZM233 | | | | | | <20 | <20 | 22 | <20 | <20 | 28 | 710 | 282 | <20 | 109 |
| | C | 2 | 16055 | | | | | | <20 | <20 | <20 | <20 | <20 | <20 | 35 | <20 | <20 | <20 |
| | 01_AE | 2 | C1080 | | | | | | <20 | <20 | <20 | <20 | <20 | <20 | 306 | <20 | <20 | <20 |
| | A1C | 2 | 246F3 | | | | | | <20 | <20 | <20 | <20 | <20 | <20 | <20 | <20 | <20 | <20 |
| | 01_AE | 2 | CM244 | | | | | | <20 | <20 | <20 | <20 | <20 | <20 | 13452 | <20 | <20 | <20 |
| | C | 2 | CE1176 | | | | | | <20 | <20 | <20 | <20 | <20 | | | | | |
| | C | 2 | CE0217 | | | | | | <20 | <20 | <20 | <20 | <20 | | | | | |
| | SIVcpzPtt | | CAM13RRK | <20 | <20 | <20 | <20 | <20 | <20 | <20 | <20 | <20 | <20 | <20 | 15579 | <20 | <20 | <20 |
| | A | 2 | SHIV.BG505 | | | | | | | | | | | | | | | |
| Viruses contaning N130 | C | 2 | CAP256_wk34 | <20 | <20 | <20 | <20 | <20 | <20 | <20 | <20 | <20 | <20 | <20 | <20 | <20 | <20 | <20 |
| | 01_AE | 2 | C4118 | | | | | | <20 | <20 | <20 | <20 | <20 | | | | | |
| | C | 2 | 1394C9 | | | | | | <20 | <20 | <20 | <20 | <20 | | | | | |
| | C | 2 | BF1266 | | | | | | <20 | <20 | <20 | <20 | <20 | | | | | |
| | C | 2 | V703_2117_110 | | | | | | <20 | <20 | <20 | <20 | <20 | | | | | |
| | C | 2 | V703_2018_240 | | | | | | <20 | <20 | <20 | <20 | <20 | | | | | |
| | C | 2 | V703_2149_060 | | | | | | <20 | <20 | <20 | <20 | <20 | | | | | |
| | A | 2 | Q168.a2 | | | | | | <20 | <20 | <20 | <20 | <20 | | | | | |
| | C | 2 | 0921.v2.c14 | | | | | | | | | | | | | | | |
| | A | 2 | 9004SS_A3_4 | | | | | | | | | | | | | | | |

| Group | Subtype | Tier | Immunogen | C1080 mRNA | | | | | BF1266-scNP | | | | | BF1266-mRNA | | | | |
|---|---|---|---|---|---|---|---|---|---|---|---|---|---|---|---|---|---|---|
| | | | NHP ID | **50** | **54** | **63** | **76** | **91** | **50** | **54** | **63** | **76** | **91** | **50** | **54** | **63** | **76** | **91** |
| | | | Study Week | **wk47** | | | | | **wk80** | | | | | **wk86** | | | | |
| | Subtype | Tier | MLV | <20 | <20 | <20 | <20 | <20 | <20 | <20 | <20 | <20 | <20 | <20 | <20 | <20 | <20 | <20 |
| Autologous Viruses | | | OPT4 | 29138 | 349406 | 234687 | 45005 | 284414 | 19172 | 358551 | 345185 | 228206 | 126678 | 6916 | 331345 | 169463 | 110461 | 97182 |
| | | | OPT4-S | 3266 | 55463 | 10131 | 5981 | 7740 | 2499 | 94607 | 55741 | 24900 | 19631 | 1042 | 37230 | 25246 | 6427 | 15135 |
| | | | OPT4-S.N160K | 95 | 424 | 4998 | 1090 | 7194 | 49 | 870 | 9434 | 5397 | 2913 | 57 | 1792 | 4439 | 3165 | 4429 |
| | | | OPT4-S.R169E | 31 | 151 | 1594 | 436 | 1602 | <20 | 2339 | 4909 | 2238 | 1803 | 63 | 894 | 2767 | 2774 | 2931 |
| Viruses lacking N130 | A | 1B | Q23.17 | <20 | 54825 | 21 | <20 | 27 | 145 | 113675 | 8826 | 1074 | 5195 | <20 | 28918 | 3224 | 224 | 2709 |
| | 02_AG | 2 | T250 | <20 | 215 | <20 | <20 | <20 | <20 | 7669 | 182 | 452 | 26 | <20 | 2203 | 54 | 111 | 81 |
| | C | 2 | ZM233 | <20 | 481 | 55 | <20 | <20 | <20 | 3174 | 5238 | 539 | 166 | <20 | 930 | 1959 | 95 | 240 |
| | C | 2 | 16055 | <20 | 493 | <20 | <20 | <20 | <20 | 3331 | 28 | <20 | 45 | <20 | 2200 | <20 | <20 | <20 |
| | 01_AE | 2 | C1080 | <20 | 18159 | <20 | <20 | <20 | 28 | 78003 | 83 | <20 | 890 | <20 | 42283 | <20 | <20 | 951 |
| | A1C | 2 | 246F3 | <20 | <20 | <20 | <20 | <20 | <20 | 117 | 40 | <20 | <20 | <20 | 36 | <20 | <20 | <20 |
| | 01_AE | 2 | CM244 | <20 | 49068 | <20 | <20 | <20 | 22 | 99206 | 705 | 47 | 475 | <20 | 37807 | 215 | <20 | 525 |
| | C | 2 | CE1176 | <20 | <20 | <20 | <20 | <20 | <20 | 138 | 0 | <20 | <20 | <20 | <20 | <20 | <20 | <20 |
| | C | 2 | CE0217 | <20 | 343 | <20 | <20 | <20 | <20 | 2311 | 181 | <20 | <20 | <20 | 513 | 26 | <20 | <20 |
| | SIVcpzPtt | | CAM13RRK | <20 | 59347 | 47 | <20 | <20 | <20 | 113960 | 3193 | 692 | 1778 | <20 | 47824 | 1057 | 318 | 580 |
| | A | 2 | SHIV.BG505 | | | | | | <20 | 3979 | 28 | <20 | 54 | <20 | 1183 | <20 | <20 | <20 |
| Viruses contaning N130 | C | 2 | CAP256_wk34 | <20 | <20 | <20 | <20 | <20 | <20 | 158 | 100 | <20 | <20 | <20 | 304 | 28 | <20 | <20 |
| | 01_AE | 2 | C4118 | <20 | <20 | <20 | <20 | <20 | <20 | 38 | <20 | <20 | <20 | <20 | 28 | <20 | <20 | <20 |
| | C | 2 | 1394C9 | <20 | <20 | <20 | <20 | <20 | <20 | 326 | 61 | <20 | <20 | <20 | 147 | 20 | <20 | <20 |
| | C | 2 | BF1266 | <20 | <20 | <20 | <20 | <20 | <20 | 95 | <20 | <20 | <20 | <20 | 595 | 1826 | <20 | 210 |
| | C | 2 | V703_2117_110 | <20 | <20 | <20 | <20 | <20 | <20 | 115 | <20 | <20 | <20 | <20 | 116 | <20 | <20 | <20 |
| | C | 2 | V703_2018_240 | <20 | <20 | <20 | <20 | <20 | <20 | 37 | 67 | <20 | <20 | <20 | <20 | <20 | <20 | <20 |
| | C | 2 | V703_2149_060 | <20 | <20 | <20 | <20 | <20 | <20 | 78 | 36 | <20 | <20 | <20 | <20 | <20 | <20 | <20 |
| | A | 2 | Q168.a2 | | | | | | <20 | 175 | <20 | <20 | <20 | <20 | 55 | <20 | <20 | <20 |
| | C | 2 | 0921.v2.c14 | | | | | | <20 | 656 | 1134 | <20 | <20 | <20 | 525 | 317 | <20 | <20 |
| | A | 2 | 9004SS_A3_4 | | | | | | <20 | 500 | 320 | 41 | 53 | <20 | 306 | 127 | <20 | <20 |

**B**

| Group | Immunogen | BF1266-scNP | | | | |
|---|---|---|---|---|---|---|
| | NHP ID | **50** | **54** | **63** | **76** | **91** |
| | Study Week | **wk80** | | | | |
| Viruses lacking N130 | Q23.17 | 69.26 | 0.1006 | 0.9795 | 11.47 | 1.488 |
| | T250 | 1324 | 1.863 | 159 | 37.29 | 476.9 |
| | ZM233 | 385.3 | 3.087 | 1.464 | 20.39 | 35.63 |
| | 16055 | >2000 | 2.619 | >2000 | >2000 | 161.4 |
| | C1080 | 342.0 | 0.0898 | 148.2 | 1604 | 5.057 |
| | 246F3 | >2000 | 123.1 | 1345 | >2000 | 1980 |
| | CM244 | 489.6 | 0.1028 | 34.59 | 370.6 | 21.04 |
| | CE1176 | >2000 | 110 | >2000 | >2000 | 1707 |
| | CE0217 | >2000 | 8.491 | 141.8 | >2000 | 530.3 |
| | CAM13RRK | >2000 | 0.0952 | 5.178 | 30.66 | 5.624 |
| | SHIV.BG505 | >2000 | 3.25 | >2000 | >2000 | 328.8 |

| Group | Immunogen | BF1266-scNP | | | | |
|---|---|---|---|---|---|---|
| | NHP ID | **50** | **54** | **63** | **76** | **91** |
| | Study Week | **wk80** | | | | |
| Viruses containing N130 | C4118 | >2000 | >2000 | >2000 | >2000 | >2000 |
| | 1394C9 | >2000 | 87.90 | 1104 | 348.3 | 518.4 |
| | BF1266 | >2000 | 455.1 | >2000 | >2000 | 1939 |
| | V703_2117_110 | >2000 | 235.1 | >2000 | >2000 | >2000 |
| | V703_2018_240 | >2000 | 667.1 | 360.0 | 528.5 | >2000 |
| | V703_2149_060 | >2000 | >2000 | 1292 | >2000 | 1469 |
| | CE0393 | 1224 | 1837 | >2000 | >2000 | 1924 |
| | 0921.v2.c14 | >2000 | 54.94 | 18.93 | 780.5 | 1263 |
| | 9004SS_A3_4 | 1321 | 121.7 | 103.4 | 531.4 | 413.4 |
| | DU123_06 | >2000 | >2000 | >2000 | >2000 | >2000 |
| | Q168.a2 | >2000 | 663.9 | >2000 | >2000 | >2000 |

**Supplemental Figure 3. Rhesus macaques serum and purified IgG for plasma neutralization ID50 titers. Related to Figure 1.**

(A) Serum neutralization titers shown as reciprocal serum dilution that inhibits 50% of virus replication. Serum was collected from each macaque over time. Gray indicates isolates that were not tested against that serum sample. (B) Neutralization titers for IgG purified from plasma shown as the concentration of IgG that inhibited 50% of virus replication (IC50) in μg/mL. The heatmaps are colored coded as green is the weakest neutralization, followed by yellow. Darker shades of red show more potent neutralization.

Supplemental Figure 4

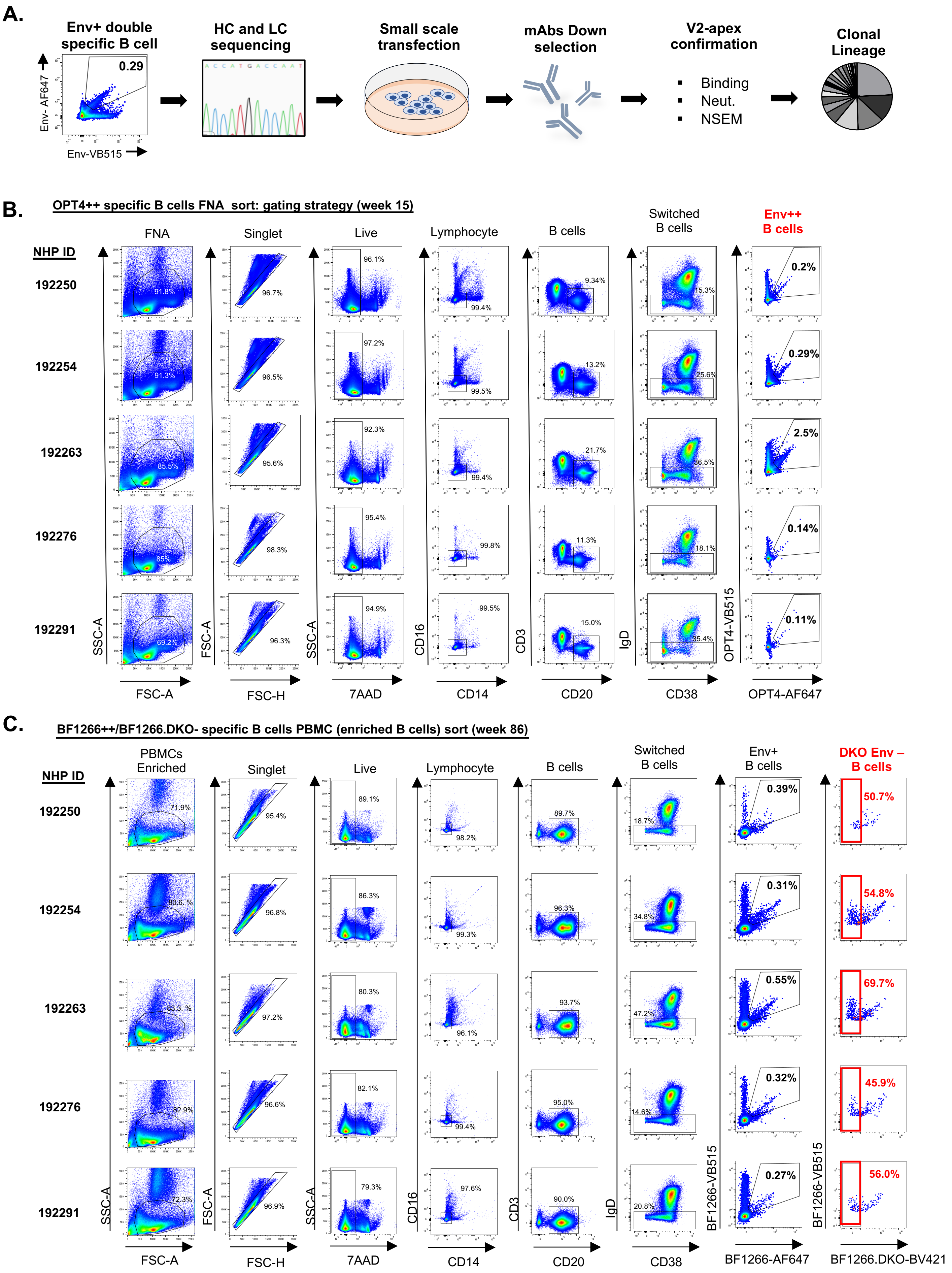

**Supplemental Figure 4. Rhesus macaques Env+ specific B cell gating strategy related to Figure 2.**

(A) Pictorial representation of the antibody discovery pipeline. (B) Gating strategy and individual dot plot of iLN FNA sort at week 15 using OPT4-labeled Env. (C). Gating strategy and individual plot of PBMCs sort at week 86 using BF1266-labeled Env.

Supplemental Figure 5

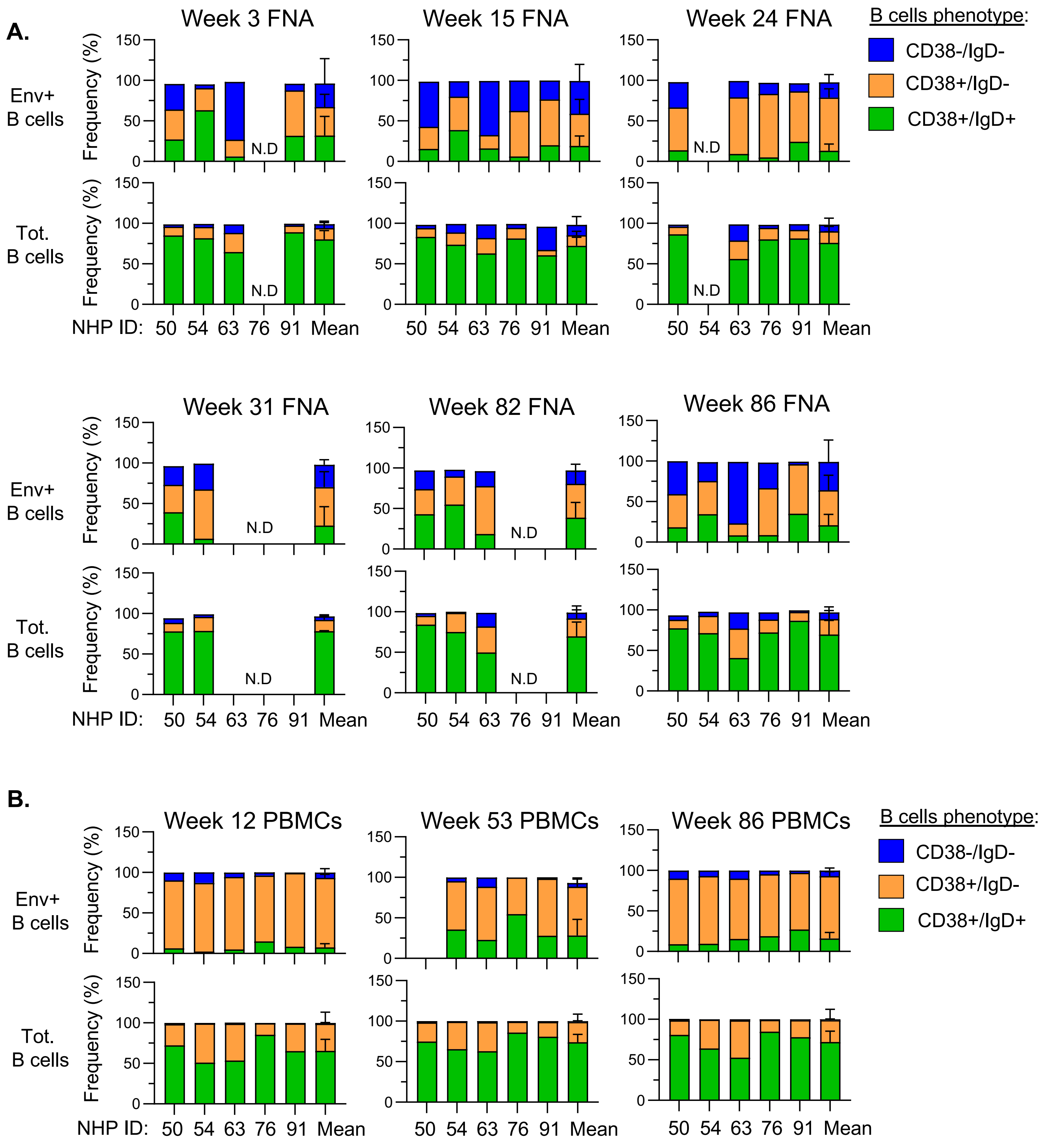


**Supplemental Figure 5. Rhesus macaques B cell phenotype related to Figure 2.**(A) Frequency of antigen specific and total B cells from isolated iLN FNA based on their CD38 and IgD expression in each NHP. (B) Frequency of antigen specific and total B cells from PBMCs based on their CD38 and IgD expression in each NHP.

Supplemental Figure 6

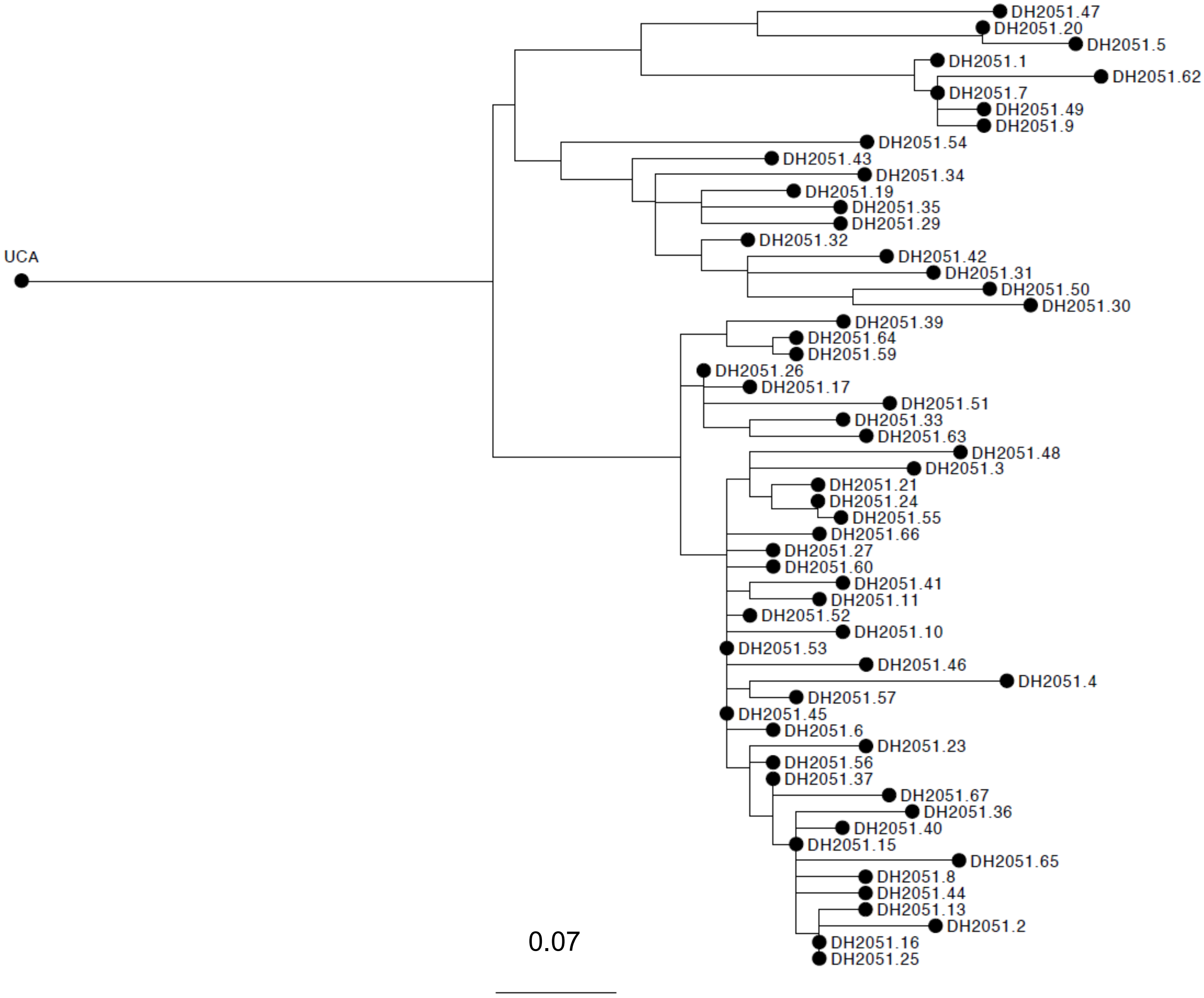


**Supplemental Figure 6. DH2051 clonal lineage related to Figure 2.** Phylogenetic Tree of 59 DH2051 clonal lineage members isolated at week 15 iLN FNA from NHP 63.

Supplemental Figure 7

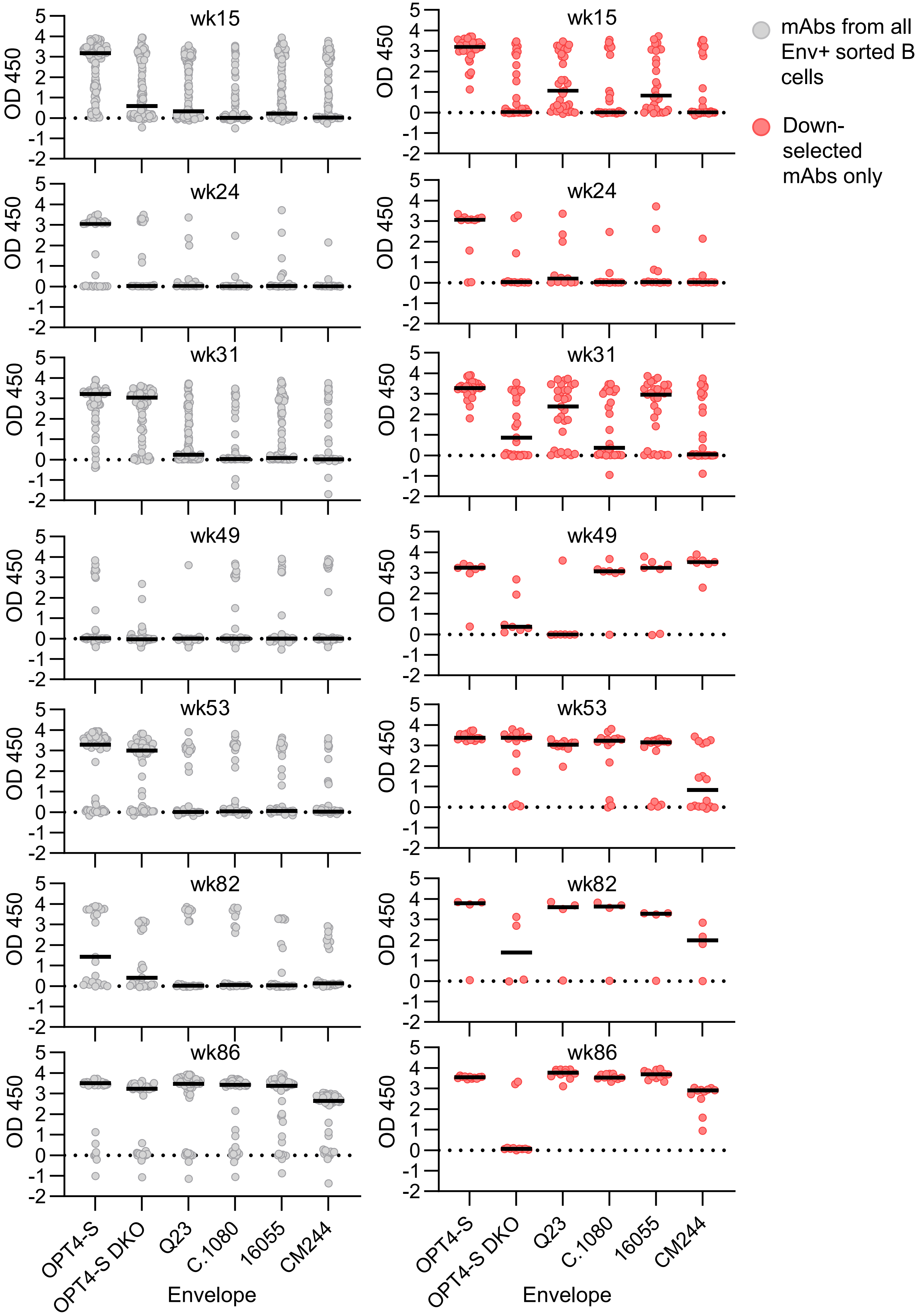

**Supplemental Figure 7. ELISA high-throughput binding screen of mAbs against a panel of HIV-1 Envs.**

Binding graphs in the left column show results from all mAbs. Binding results in the right column show binding of mAbs selected for further study, based on genotype or phenotype. Antibodies are separated by time point from which they were isolated. Horizontal bars show median binding value for each envelope. OPT4-S DKO, OPT4-S_N160K_R169E.

### Supplemental Figure 8

| NHP ID | Study week | mAbs ID | OPT4 | OPT4-S | OPT4-S R169E | OPT4-S _N160K | Q23.17 | CAM13RRK | CM244 | C1080 | 16055 | CAP256.wk34 |
|---|---|---|---|---|---|---|---|---|---|---|---|---|
| 50 | 15 | DH2053 | >300 | 30.55 | >300 | >300 | >300 | >300 | >300 | >300 | >300 | 298.40 |
| 50 | 15 | DH2054.1 | >300 | 60.84 | >300 | >300 | >300 | >300 | >300 | >300 | >300 | >300 |
| 50 | 15 | DH2054.2 | >300 | 140.10 | >300 | >300 | >300 | >300 | >300 | >300 | >300 | >300 |
| 50 | 15 | DH2056.1 | >300 | 107.20 | >300 | 272.30 | >300 | >300 | >300 | >300 | >300 | >300 |
| 50 | 15 | DH2058 | 11.03 | 1.78 | >300 | 234.30 | >300 | >300 | >300 | >300 | >300 | >300 |
| 50 | 31 | DH2056.1 | >300 | 19.22 | >300 | >300 | >300 | >300 | >300 | >300 | >300 | >300 |
| 50 | 31 | DH2059.1 | 0.05 | 0.12 | >300 | >300 | >300 | >300 | >300 | 166.10 | >300 | >300 |
| 50 | 31 | DH2060 | 0.01 | 0.01 | 163.30 | >300 | 161.60 | 1.82 | 243.00 | 164.20 | 251.00 | >300 |
| 50 | 31 | DH2061.1 | 0.54 | 0.79 | >300 | >300 | >300 | 179.70 | 290.80 | >300 | >300 | >300 |
| 50 | 31 | DH2061.2 | 0.38 | 0.32 | >300 | >300 | >300 | >300 | >300 | >300 | >300 | >300 |
| 50 | 31 | DH2061.3 | 0.05 | 0.13 | >300 | >300 | >300 | 221.50 | >300 | >300 | >300 | >300 |
| 50 | 31 | DH2062.1 | 35.50 | 4.35 | 243.60 | >300 | >300 | 294.60 | >300 | >300 | >300 | >300 |
| 50 | 31 | DH2062.2 | >300 | 1.52 | 127.40 | >300 | >300 | >300 | >300 | >300 | >300 | >300 |
| 50 | 31 | DH2063 | 2.32 | 8.35 | >300 | >300 | >300 | 290.30 | >300 | >300 | >300 | >300 |
| 50 | 31 | DH2065 | 77.81 | 3.17 | 270.60 | >300 | >300 | 181.80 | >300 | >300 | 291.90 | >300 |
| 50 | 31 | DH2066 | 203.50 | >300 | 134.10 | >300 | >300 | 279.90 | >300 | 254.80 | >300 | >300 |
| 50 | 31 | DH2068 | 152.90 | 117.10 | 161.40 | >300 | >300 | 132.00 | 273.80 | 241.60 | 276.40 | >300 |
| 50 | 82 | DH2059.2 | 0.01 | 0.17 | >300 | >300 | 201.40 | >300 | >300 | >300 | >300 | >300 |
| 50 | 82 | DH2059.3 | 0.04 | 0.39 | >300 | >300 | >300 | >300 | >300 | >300 | >300 | >300 |
| 50 | 82 | DH2061.4 | 0.05 | 0.16 | >300 | >300 | >300 | >300 | >300 | >300 | >300 | >300 |
| 54 | 15 | DH2050.1 | 0.01 | 0.01 | 221.50 | 294.40 | 0.01 | 3.72 | 12.38 | 288.60 | 289.70 | >300 |
| 54 | 15 | DH2050.2 | 0.34 | 0.48 | >300 | >300 | 29.85 | 39.33 | 177.10 | >300 | >300 | >300 |
| 54 | 15 | DH2050.3 | 0.01 | 0.68 | >300 | >300 | 0.85 | >300 | >300 | >300 | >300 | >300 |
| 54 | 15 | DH2050.4 | 0.01 | 0.05 | >300 | >300 | 2.14 | 227.10 | 279.90 | >300 | >300 | >300 |
| 54 | 15 | DH2071 | 110.20 | 132.60 | >300 | >300 | >300 | >300 | >300 | >300 | >300 | >300 |
| 54 | 15 | DH2072.1 | >300 | >300 | >300 | >300 | >300 | >300 | >300 | >300 | >300 | >300 |
| 54 | 15 | DH2072.2 | >300 | 5.16 | >300 | >300 | >300 | >300 | >300 | >300 | >300 | >300 |
| 54 | 15 | DH2072.3 | >300 | 153.70 | >300 | >300 | >300 | >300 | >300 | >300 | >300 | >300 |
| 54 | 15 | DH2073.1 | 280.30 | 33.51 | >300 | >300 | >300 | >300 | >300 | >300 | >300 | >300 |
| 54 | 15 | DH2074 | 224.40 | 28.54 | >300 | >300 | >300 | 290.80 | >300 | >300 | >300 | >300 |
| 54 | 15 | DH2075 | 4.99 | 0.30 | >300 | >300 | >300 | >300 | >300 | >300 | >300 | >300 |
| 54 | 15 | DH2076 | >300 | >300 | >300 | >300 | >300 | >300 | >300 | >300 | >300 | >300 |
| 54 | 15 | DH2077 | 1.36 | 2.30 | >300 | >300 | >300 | >300 | >300 | >300 | >300 | >300 |
| 54 | 15 | DH2078.1 | >300 | 4.74 | >300 | >300 | >300 | >300 | >300 | >300 | >300 | >300 |
| 54 | 15 | DH2081 | 176.00 | 34.28 | >300 | >300 | >300 | >300 | >300 | >300 | 240.80 | >300 |
| 54 | 31 | DH2050.10 | 0.01 | 0.09 | >300 | >300 | 49.09 | >300 | >300 | >300 | >300 | >300 |
| 54 | 31 | DH2050.5 | 0.01 | 0.01 | >300 | >300 | 0.01 | 1.57 | 0.33 | >300 | >300 | >300 |
| 54 | 31 | DH2050.6 | 0.01 | 0.01 | >300 | 252.90 | 0.01 | 0.49 | 0.26 | 74.21 | >300 | >300 |
| 54 | 31 | DH2050.7 | 0.21 | 1.61 | >300 | >300 | 17.64 | 34.59 | 91.10 | >300 | >300 | >300 |
| 54 | 31 | DH2050.8 | 0.01 | 0.01 | >300 | >300 | 0.01 | 0.01 | 0.01 | 61.23 | 144.90 | >300 |
| 54 | 31 | DH2050.9 | 0.01 | 0.01 | >300 | >300 | 0.01 | 0.88 | 3.07 | 176.00 | >300 | 294.30 |
| 54 | 31 | DH2072.4 | >300 | 3.27 | >300 | >300 | >300 | >300 | >300 | >300 | >300 | >300 |
| 54 | 31 | DH2072.5 | >300 | >300 | >300 | >300 | >300 | >300 | >300 | >300 | >300 | >300 |
| 54 | 31 | DH2072.6 | >300 | 8.15 | >300 | >300 | >300 | >300 | >300 | >300 | >300 | >300 |
| 54 | 31 | DH2072.7 | 268.70 | >300 | >300 | >300 | >300 | >300 | >300 | >300 | >300 | >300 |
| 54 | 31 | DH2073.2 | 0.01 | 0.08 | >300 | >300 | >300 | >300 | >300 | >300 | >300 | 286.00 |
| 54 | 31 | DH2079 | >300 | 63.34 | >300 | >300 | >300 | >300 | >300 | >300 | >300 | 256.70 |
| 54 | 31 | DH2080 | >300 | 15.72 | >300 | >300 | >300 | >300 | >300 | >300 | >300 | >300 |
| 54 | 31 | DH2082 | >300 | 188.60 | >300 | >300 | >300 | >300 | >300 | >300 | >300 | >300 |
| 54 | 31 | DH2083 | >300 | 60.83 | >300 | >300 | >300 | >300 | >300 | >300 | >300 | >300 |
| 54 | 53 | DH2050.11 | 0.01 | 0.02 | 47.18 | >300 | 0.01 | 0.01 | 0.01 | 0.01 | 0.20 | 149.60 |
| 54 | 82 | DH2050.12 | 0.01 | 0.01 | >300 | >300 | 0.01 | 0.37 | 0.01 | 0.19 | >300 | >300 |
| 54 | 82 | DH2085 | >300 | >300 | >300 | >300 | >300 | >300 | >300 | >300 | >300 | >300 |
| 54 | 86 | DH2050.13 | 0.01 | 0.01 | >300 | >300 | 0.01 | 0.01 | 0.01 | 0.25 | 0.08 | 236.70 |
| 54 | 86 | DH2050.14 | 0.01 | 0.01 | >300 | >300 | 0.01 | 0.01 | 0.01 | 0.01 | 0.02 | 29.39 |
| 54 | 86 | DH2050.15 | 0.01 | 0.01 | >300 | >300 | 0.01 | 0.01 | 0.01 | 0.01 | 0.30 | 0.14 |
| 54 | 86 | DH2050.16 | 0.01 | 0.01 | >300 | >300 | 0.01 | 0.01 | 0.01 | 0.01 | 0.04 | 0.08 |
| 54 | 86 | DH2050.17 | 0.01 | 0.01 | >300 | >300 | 0.01 | 0.01 | 0.01 | 0.01 | 0.08 | 0.20 |
| 54 | 86 | DH2050.18 | 0.01 | 0.01 | >300 | >300 | 0.01 | 0.01 | 0.01 | 0.20 | 73.45 | 0.20 |
| 54 | 86 | DH2050.19 | 0.01 | 0.01 | >300 | >300 | 0.01 | 0.01 | 0.01 | 0.75 | 0.56 | 0.29 |
| 63 | 15 | DH2051.1 | 0.01 | 0.01 | >300 | 278.50 | 0.02 | 0.01 | 133.80 | >300 | >300 | >300 |
| 63 | 15 | DH2051.2 | 0.01 | 0.44 | >300 | >300 | >300 | 211.20 | >300 | >300 | >300 | >300 |
| 63 | 15 | DH2051.3 | 0.08 | 7.37 | >300 | >300 | >300 | >300 | >300 | >300 | >300 | >300 |
| 63 | 15 | DH2051.4 | 0.01 | 0.03 | >300 | >300 | >300 | 0.57 | >300 | >300 | >300 | >300 |
| 63 | 15 | DH2051.5 | 0.01 | 0.01 | >300 | >300 | 41.35 | 0.05 | >300 | >300 | >300 | >300 |
| 63 | 15 | DH2051.6 | 0.01 | 0.02 | >300 | >300 | 2.77 | 1.27 | >300 | >300 | >300 | >300 |
| 63 | 15 | DH2051.7 | 0.01 | 0.01 | >300 | >300 | 0.04 | 0.02 | 257.60 | >300 | >300 | >300 |
| 63 | 15 | DH2087 | 2.16 | 0.50 | >300 | >300 | >300 | >300 | >300 | >300 | >300 | >300 |
| 63 | 15 | DH2089 | 140.30 | 29.67 | >300 | >300 | >300 | >300 | >300 | >300 | >300 | 224.10 |
| 63 | 24 | DH2088 | 38.34 | 8.65 | >300 | >300 | >300 | 160.00 | >300 | >300 | 281.20 | >300 |
| 63 | 24 | DH2090.1 | 2.28 | 1.71 | >300 | >300 | >300 | >300 | >300 | >300 | >300 | >300 |
| 63 | 53 | DH2094 | 0.01 | 0.01 | >300 | >300 | >300 | >300 | >300 | >300 | >300 | >300 |
| 63 | 86 | DH2090.2 | 0.01 | 0.01 | >300 | >300 | 11.44 | 1.52 | >300 | >300 | >300 | >300 |
| 63 | 86 | DH2090.3 | 0.01 | 0.01 | >300 | >300 | >300 | >300 | >300 | >300 | >300 | >300 |
| 63 | 86 | DH2095 | 0.01 | 0.01 | >300 | >300 | 0.24 | 0.21 | 1.85 | >300 | >300 | 245.80 |
| 76 | 15 | DH2097.1 | 278.60 | 109.70 | 296.60 | >300 | >300 | 226.00 | >300 | >300 | 271.40 | >300 |
| 91 | 15 | DH2099 | >300 | >300 | >300 | >300 | >300 | >300 | >300 | >300 | >300 | >300 |
| 91 | 24 | DH2101.3 | 173.00 | 8.31 | 231.90 | >300 | >300 | 176.00 | 279.60 | 233.00 | 215.50 | >300 |
| 91 | 24 | DH2103 | 40.92 | 3.85 | >300 | >300 | >300 | 296.70 | >300 | >300 | >300 | >300 |
| 91 | 24 | DH2111 | >300 | >300 | >300 | >300 | >300 | >300 | >300 | >300 | >300 | >300 |
| 91 | 24 | DH2112 | 0.22 | 0.27 | >300 | >300 | >300 | >300 | >300 | >300 | >300 | >300 |
| 91 | 53 | DH2100 | >300 | 21.75 | >300 | >300 | >300 | >300 | >300 | >300 | >300 | >300 |
| 91 | 53 | DH2101.4 | 0.02 | 0.08 | >300 | >300 | >300 | >300 | >300 | >300 | >300 | >300 |
| 91 | 53 | DH2102.1 | 0.01 | 0.02 | >300 | >300 | >300 | 295.20 | >300 | >300 | >300 | >300 |
| 91 | 53 | DH2102.2 | 0.01 | 0.04 | >300 | 246.50 | >300 | >300 | >300 | >300 | >300 | >300 |
| 91 | 53 | DH2106.6 | 0.01 | 0.01 | >300 | >300 | 6.32 | 0.23 | >300 | >300 | 267.20 | >300 |
| 91 | 53 | DH2106.7 | 0.01 | 0.01 | >300 | >300 | 59.06 | 21.96 | >300 | >300 | 43.42 | >300 |
| 91 | 53 | DH2107 | 0.04 | 0.09 | >300 | >300 | >300 | >300 | >300 | >300 | >300 | >300 |
| 91 | 53 | DH2108 | 0.01 | 0.01 | >300 | >300 | >300 | >300 | >300 | >300 | >300 | >300 |
| 91 | 53 | DH2109 | 0.01 | 0.02 | >300 | 52.96 | >300 | >300 | >300 | >300 | >300 | >300 |
| 91 | 86 | DH2101.5 | 0.03 | 0.07 | >300 | >300 | 0.34 | >300 | >300 | >300 | >300 | >300 |

IC50 (µg/mL)
>300
<300
<50
<10
<5
<1
<0.05

**Supplemental Figure 8. IC50 neutralization of the V2-apex mAbs down selected related to Figure 2**

Neutralization IC50 table of the 91 down selected V2-apex mAbs against a panel of HIV-1.

**Supplemental Figure 9**

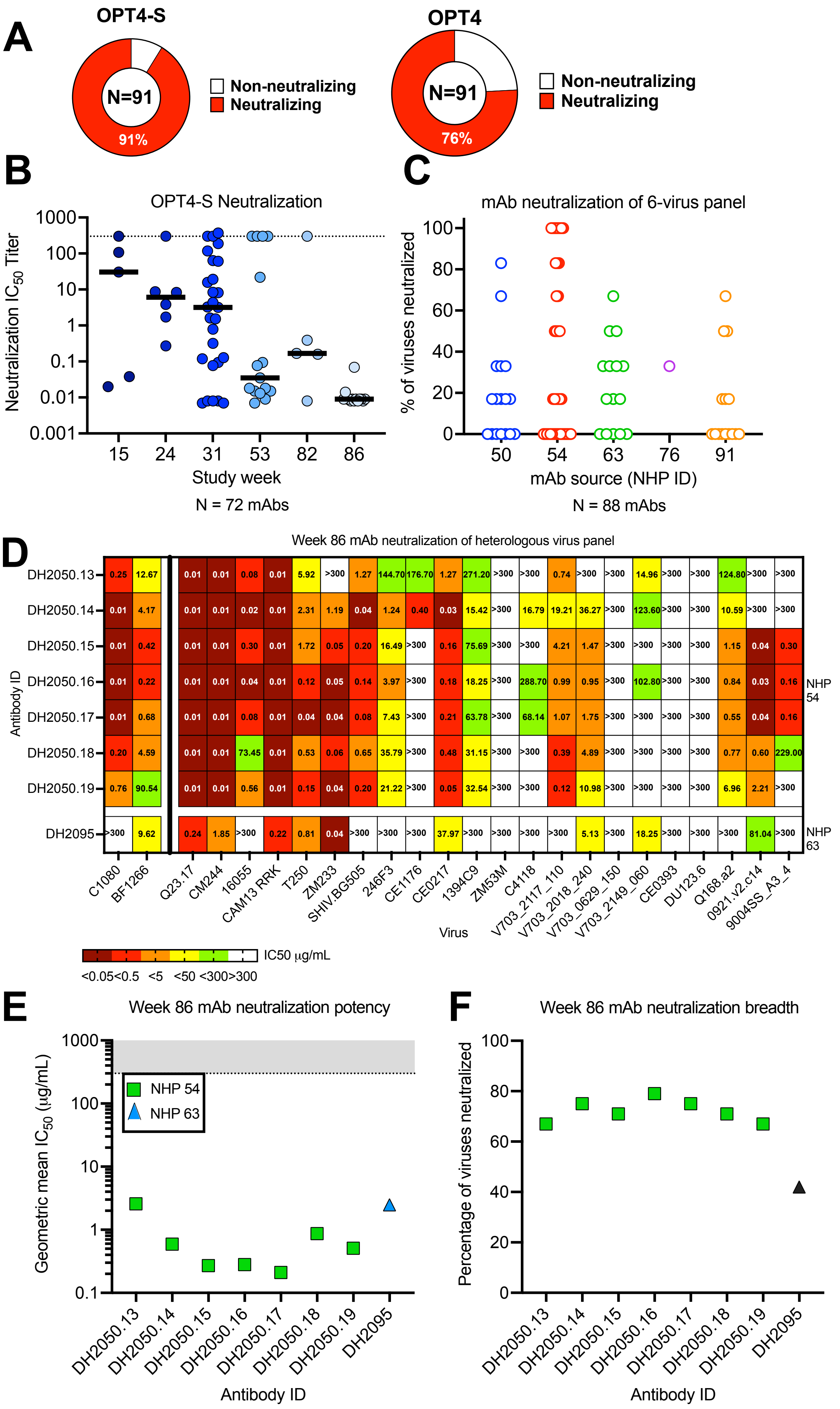

**Supplemental Figure 9. Summary of neutralization activity of monoclonal antibodies isolated at different timepoints from each NHP related to Figure 2.** **(**A) The percentage of characterize monoclonal antibodies (N=91) that neutralized OPT4-S or OPT4 viruses. (B) Neutralization potency (IC50 in μg/mL) of the antibodies against OPT4-S over time. Horizontal bar shows geometric mean for all antibodies at the time point. (C) Neutralization breadth (%) for 88 down-selected mAbs against Q23.17, C.1080. CAM13.RRK, CAP256.34.c80, CM244, and 16055. (D) Neutralization IC50 titer (in μg/mL) for week 86 mAbs from NHPs 54 and 63 against a panel of 24 viruses. (E-D) Neutralization potency and breadth against the 24-virus panel for week 86 mAbs.

Supplemental Figure 10

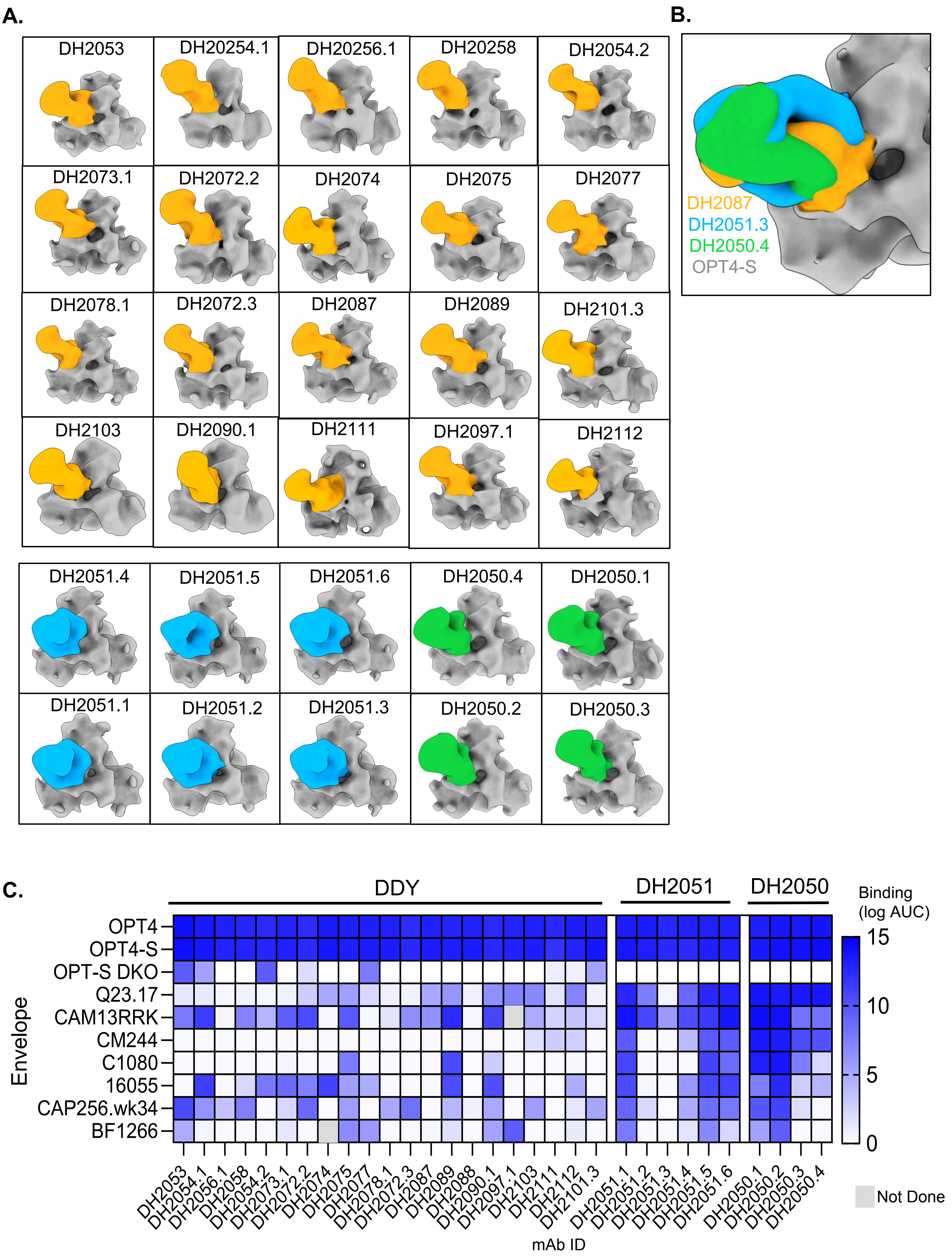

**Supplemental Figure 10. OPT4-scNP induced V2-apex mAbs NSEM and binding profile related to Figure 3.**

(A) NSEM of isolated V2-apex mAbs: DYY motifs (yellow); DH2051 lineage (Blue) and DH2050 lineage (green) complexed with OPT4-S soluble trimers. (B) NSEM merge of a representative of isolated V2-apex mAbs. (C) Isolated V2-apex mAbs binding by ELISA against a panel of soluble envelopes trimers

Supplemental Figure 11

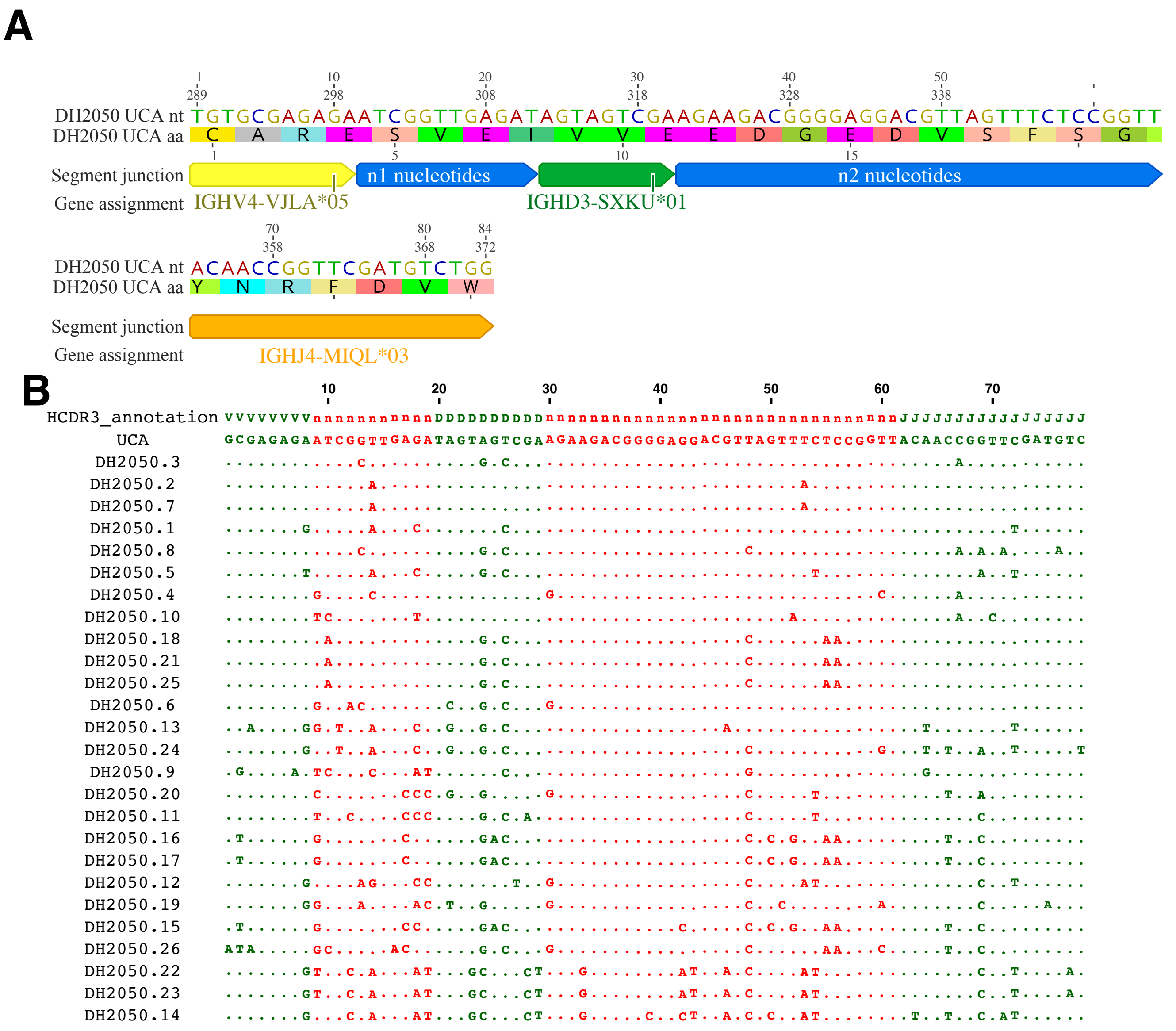

**Supplemental Figure 11. DH2050 HCDR3 junctional analysis showing the EDGED motif positioned in the n2 region of the HCDR3. (A)** HCDR3 junctional analysis, which partitions the HCDR3 by gene segments and n1 and n2 regions. Both nucleotide and amino acid sequences are shown for the inferred precursor of the DH2050 lineage (called the unmutated common ancestor, UCA). Gene segment assignments are shown on the bottom row. **(B)** Nucleotide somatic mutations are shown for each HCDR3 region. Each row shows the sequence of a different DH2050 clone member. The top row denotes HCDR3 position as variable (V), diversity (D), joining (J), or n1 or n2 (n). Periods indicate identical nucleotide compared to the UCA.

Supplemental Figure 12

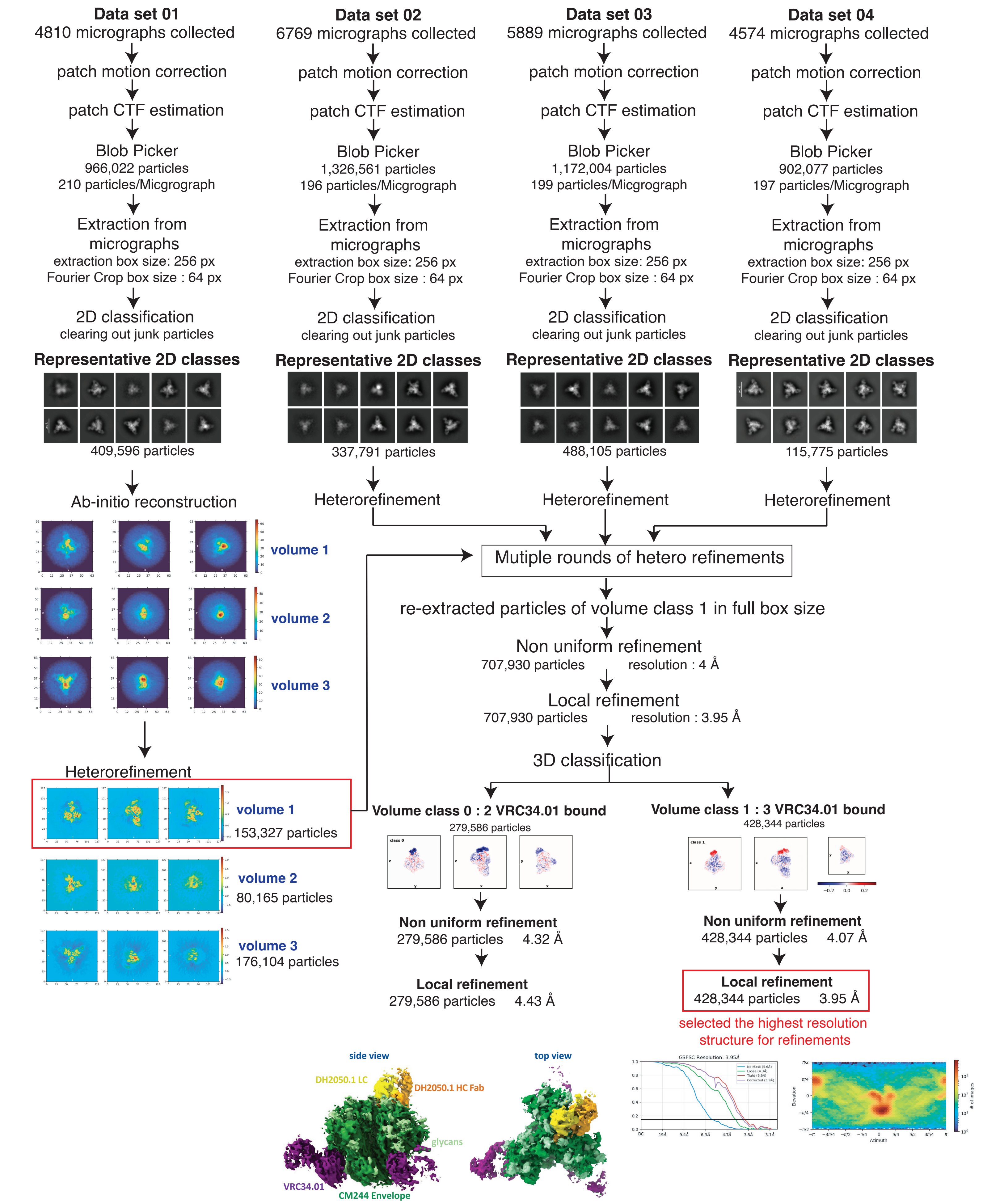


**Supplemental Figure 12. CryoSparc data processing workflow for the CM244 envelope in complex with DH2050.1 Fab.**

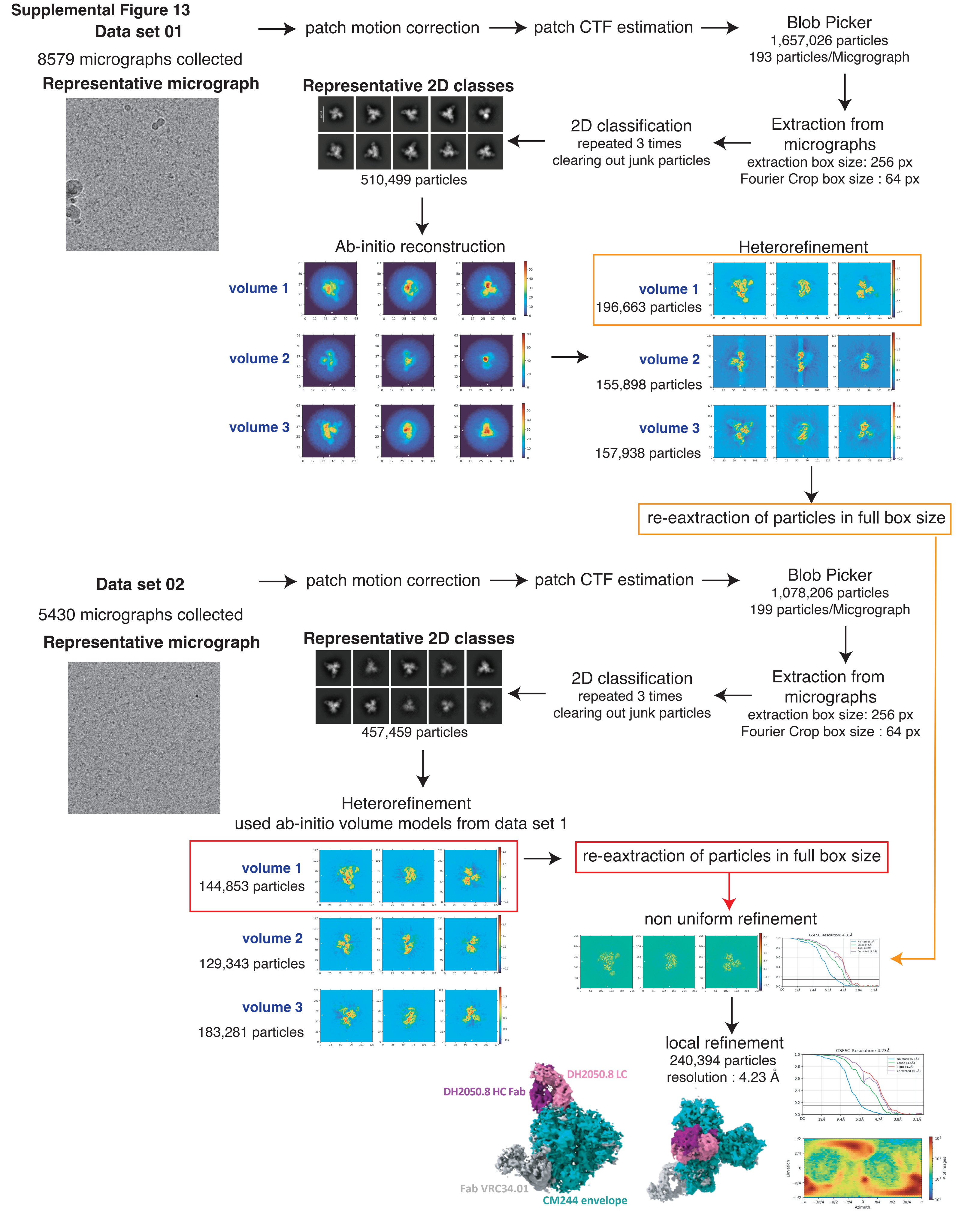


**Supplemental Figure 13. CryoSparc data processing workflow for the CM244 envelope in complex with DH2050.8 Fab.**

Supplemental Figure 14

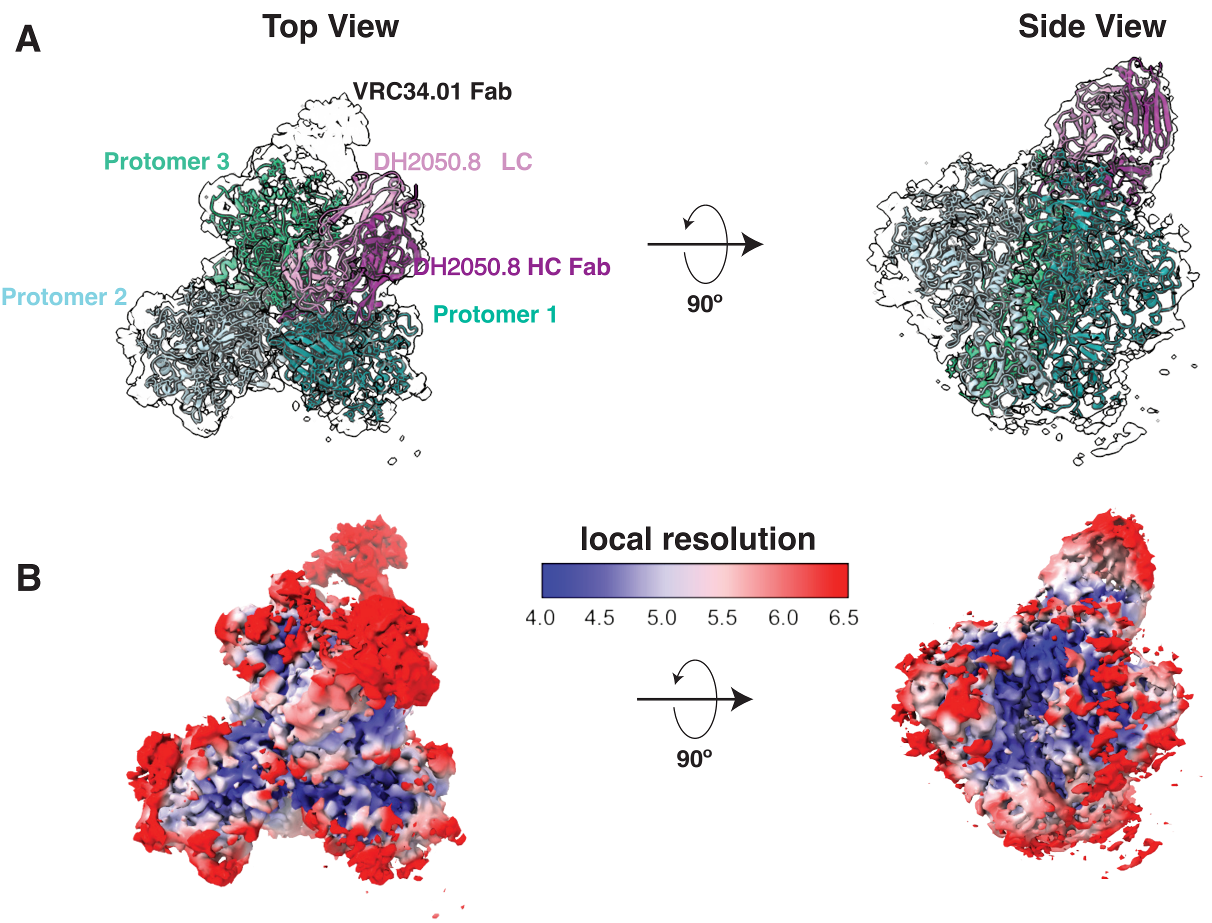


**Supplemental Figure 14. Cryo-EM model fitting for DH2050.8. (A)** The model fitting of the CM244 envelope and DH2050.8 Fab to the Cryo-EM map. **(B)** The local resolution of the Cryo-EM maps of the CM244 envelope in complex with DH2050.8 Fab

Supplemental Figure 15

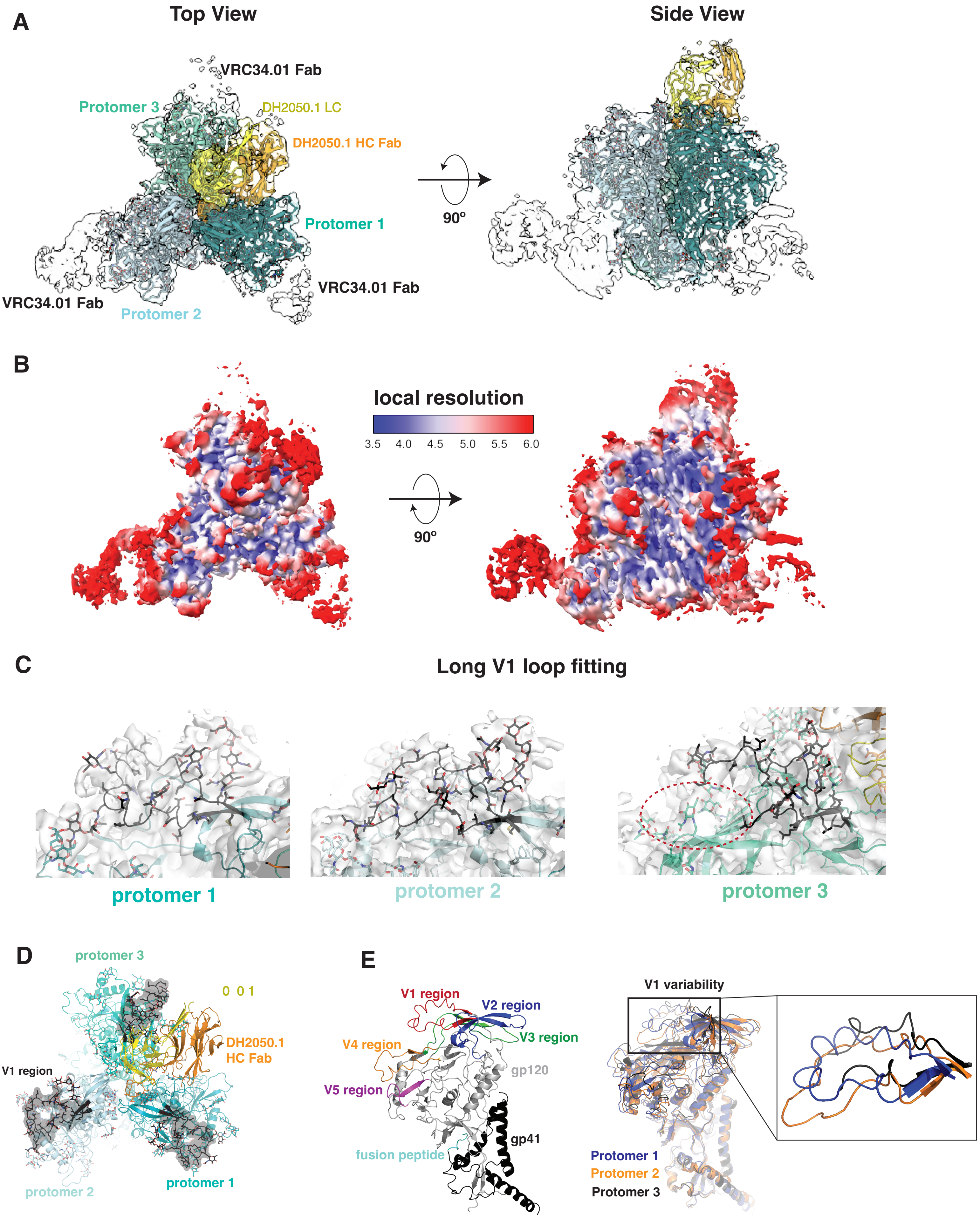


**Supplemental Figure 15. Cryo-EM maps of the CM244 envelope in complex with DH2050.1 Fab. (A)** The model fitting of the CM244 envelope and DH2050.1 Fab to the Cryo-EM map. **(B)** The local resolution of the Cryo-EM maps of the CM244 envelope in complex with DH2050.1 Fab. **(C)** Fitting of the long V1 loop, which is characteristic of the CM244 envelope, to the Cryo-EM map. **(D)** Position of the long V1 loop (colored in black highlighted with the surface) with respect to the bound antibody DH2050.1. **(E)** The alignment of the three protomers of the CM244 envelope highlights the variability of the V1 loop between the protomers.

Supplemental Figure 16

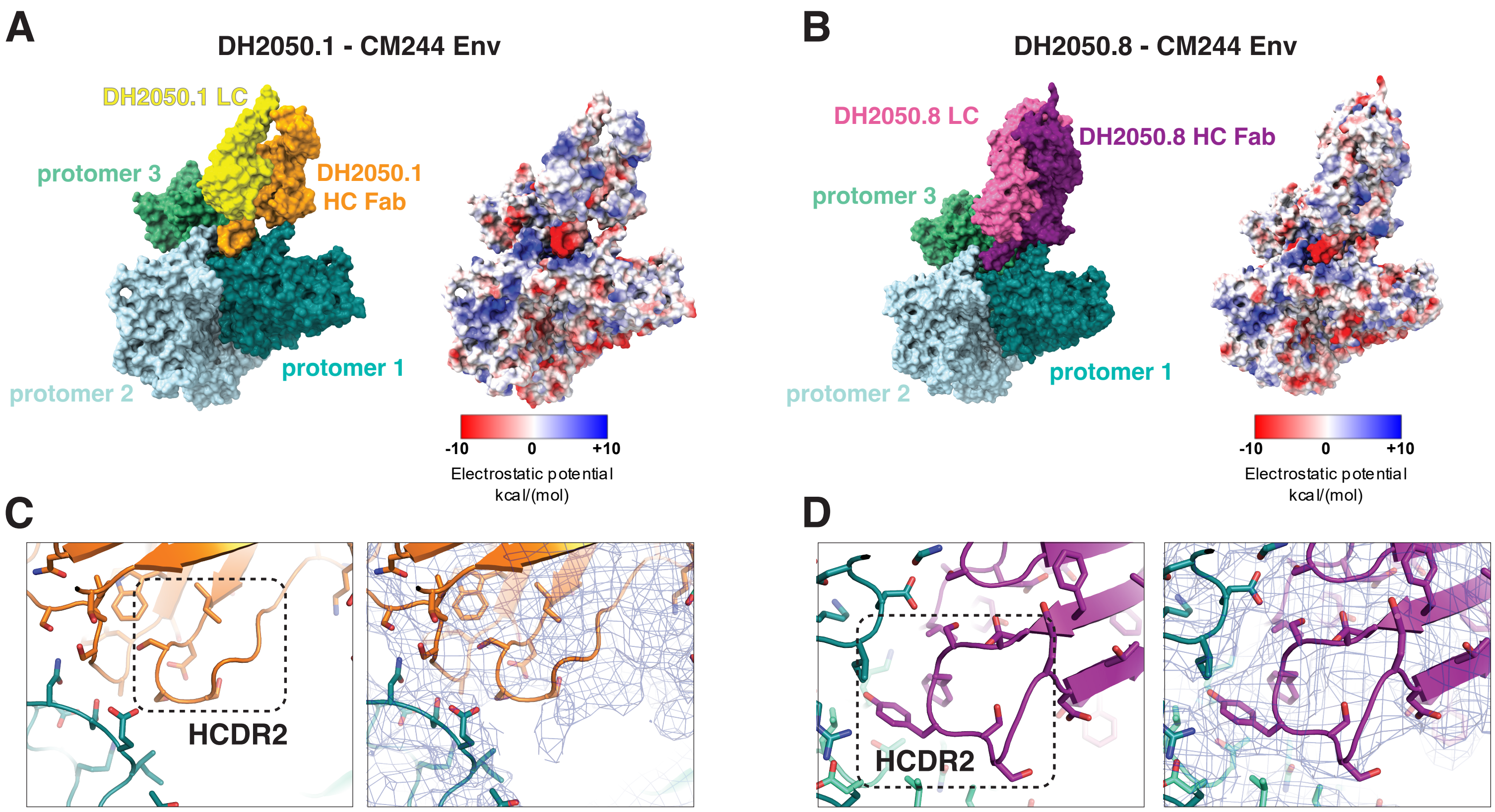


**Supplemental Figure 16. HCDR3 electrostatic potential. (A)** The electrostatic potential distribution of the structure of CM244 envelope in complex with DH2050.1 Fab. **(B)** The electrostatic potential distribution of the structure of CM244 envelope in complex with DH2050.8 Fab. **(C)** The HCDR2 loop of the DH2050.1 Fab and the model fitting to the Cryo-EM map. **(D)** The HCDR2 loop of the DH2050.8 Fab and the model fitting to the Cryo-EM map.

Supplemental Figure 17

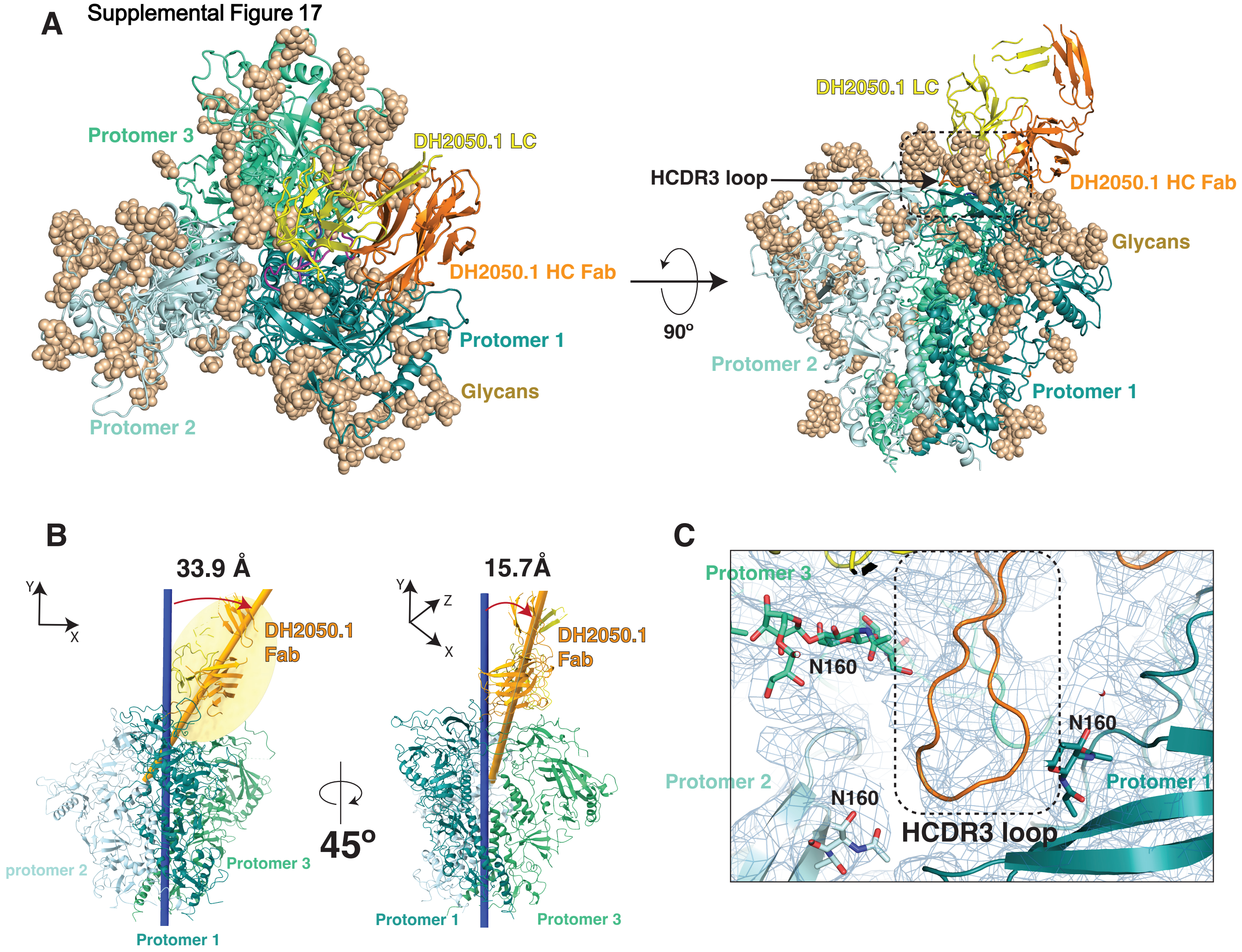

**Supplemental Figure 17. DH2050.1 binding to CM244 Env trimer. (A) Top and side view of DH2050.1 binding to CM244 Env. Glycans are shown in sphere represenation (tan color). (B)** Approach angle of DH2050.1 relative to the Env central axis. **(C)** The HCDR3 loop and N160 glycan model fitting to the Cryo-EM map showing interactions between the HCDR3 loop and N160 glycan.

Supplemental Figure 18

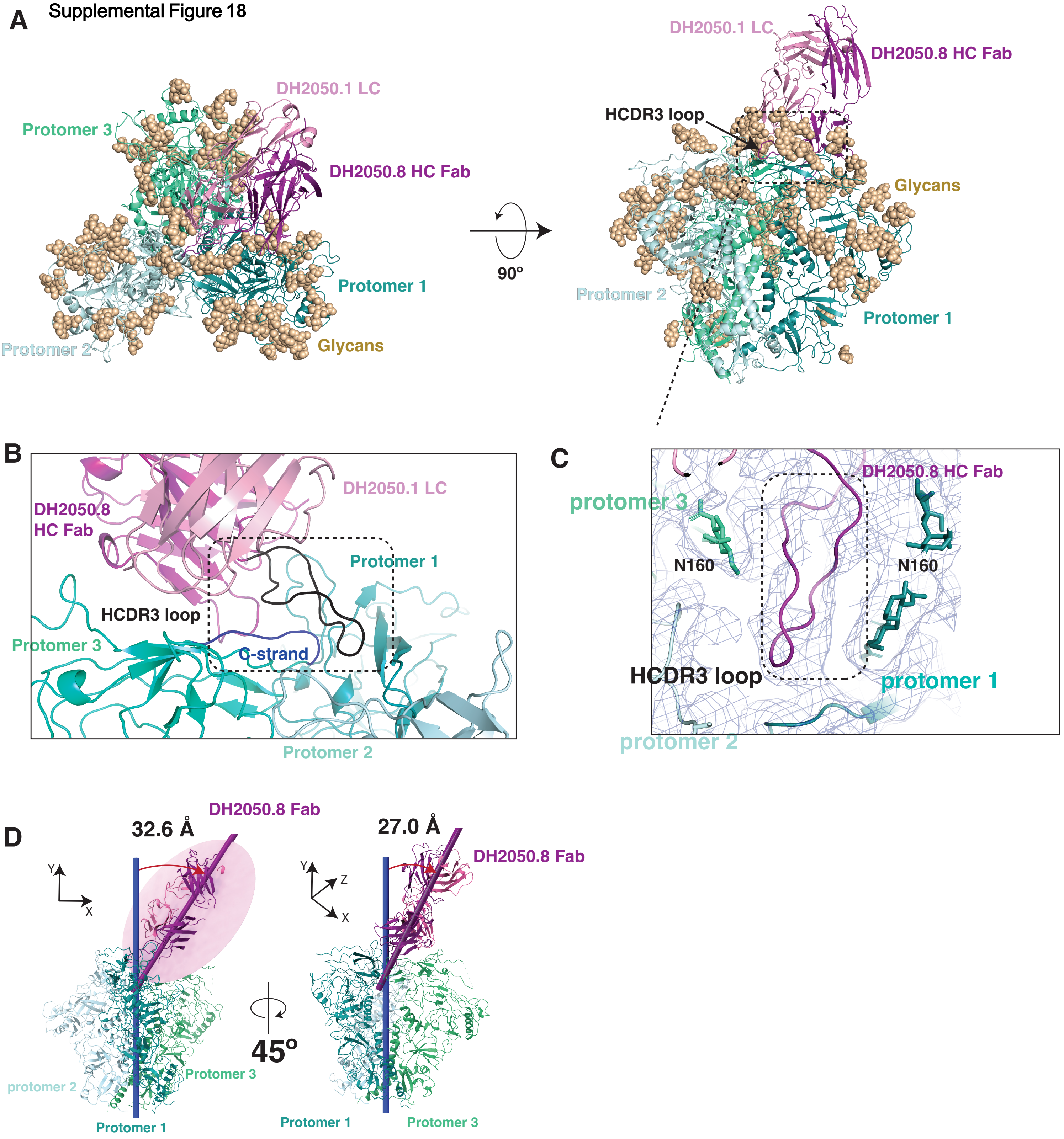

**Supplemental Figure 18. DH2050.8 binding to CM244 Env trimer.** (**A**) Top and side view of DH2050.8 binding to CM244 Env. Glycans are shown in sphere represenation (tan color). (**B**) DH2050.8 HCDR3 position relative to the Env V2 Apex region. (**C**) The HCDR3 loop and N160 glycan model fitting to the Cryo-EM map showing interactions between the HCDR3 loop and N160 glycan. (**D**) Approach angle of DH2050.8 relative to the Env central axis.
